# JBrowse 2: A modular genome browser with views of synteny and structural variation

**DOI:** 10.1101/2022.07.28.501447

**Authors:** Colin Diesh, Garrett J Stevens, Peter Xie, Teresa De Jesus Martinez, Elliot A. Hershberg, Angel Leung, Emma Guo, Shihab Dider, Junjun Zhang, Caroline Bridge, Gregory Hogue, Andrew Duncan, Matthew Morgan, Tia Flores, Benjamin N. Bimber, Robin Haw, Scott Cain, Robert M. Buels, Lincoln D. Stein, Ian H. Holmes

## Abstract

We present JBrowse 2, a general-purpose genome annotation browser offering enhanced visualization of complex structural variation and evolutionary relationships. JBrowse 2 retains the core features of the open-source JavaScript genome browser JBrowse while adding new views for synteny, dotplots, breakpoints, gene fusions, and whole-genome overviews. The software readily allows users to share sessions, open multiple genomes or views, and navigate quickly between these views. It can be embedded in a web page, used as a standalone desktop application, or run from Jupyter notebooks or R sessions. Using a plugin framework, developers can create new data adapters, track types, and visualizations. These improvements are enabled by a ground-up redesign of the JBrowse architecture using modern web technology. We describe application functionality, use cases, performance benchmarks, and implementation notes for web administrators and developers.

## Background

Genome browsers are a fundamental visualization and analysis tool for genomics. As the technology underpinning the field has progressed—from the study of individual genes, through whole genomes, up to multiple related genomes—the linear DNA sequence has provided a natural visual frame for presenting biological hypotheses (such as annotated gene and variant locations) alongside the primary evidence for those hypotheses. While the genome browser has proved long-lived as a visualization tool, the progression of sequencing technology has influenced the types of visualization needed. Sequencing is now sufficiently affordable that population genomics and comparative genomics have become commonplace. Long-read sequencing has enabled the investigation of structural variation, resolution of individual genotypes from a mixture, long haplotypes, and improved genome assemblies that were inaccessible using short reads. In addition, a diversity of sequencing kits are generating a wealth of data on epigenetic and transient states of the cell, such as DNA-protein associations, methylation, and RNA transcript levels. All of this information is genome-mappable and therefore viewable in a genome browser, but the new technology demands new visualization tools—and modalities— to represent it appropriately.

### History of JBrowse and the GMOD project

The tools and resources maintained by the Generic Model Organism Database project (GMOD) have enabled many genome projects to develop their own genome databases and websites. The GMOD project has developed and maintained two genome browsers: i) the Perl-based GBrowse genome annotation browser (Stein et al. 2002), the first portable web-based genome browser to achieve widespread adoption; and ii) the JavaScript browser JBrowse (Skinner et al. 2009; Buels et al. 2016), which introduced client-side rendering, a single-page user interface that avoided page reloads, drag-and-drop annotation tracks, animated panning and zooming transitions, and a static-site deployment model.

The original JBrowse app (henceforth “JBrowse 1”) has been reliable and extremely popular. However, it has become increasingly difficult to extend JBrowse 1 due to deep-rooted design assumptions (such as the assumption that only one genome would ever be displayed) and its dependence on older software libraries. This paper describes JBrowse 2, a complete rewrite of JBrowse 1 with a similar user interface but a modern software architecture. As we report in this manuscript, JBrowse 2 goes well beyond the capabilities of JBrowse 1. JBrowse 2 is particularly well-suited to visualizing genomic structural variants and evolutionary relationships among genes and genomes with syntenic visualizations.

### Structural variant and synteny visualization tools

Many genome browsers, including the GMOD browsers listed above, use a reference genome to provide a coordinate system in which to align annotations and evidence, including related genomes. This paradigm is ideal for visualizing individual reference genomes and small localized variants, such as single nucleotide polymorphisms and small indels. However, within this paradigm, it can become complicated to visualize structural variations whose alignment to the reference coordinate system departs strongly from collinearity, such as big duplications, deletions, translocations, insertions, inversions and other complex rearrangements. A similar point holds regarding visualization of synteny between genomes: inter-genome alignments can be collinear over small scales but structurally disrupted over larger scales.

Several specialized tools have been developed to visualize synteny or structural variation. GBrowse-Syn is an interactive tool that allows comparison of regions of multiple genomes against a reference sequence (McKay, Vergara, and Stajich 2010). It uses a joining database representing links between different species and maps sequence coordinates in the aligned segments or synteny blocks (McKay, Vergara, and Stajich 2010). The Artemis Comparison Tool (ACT) enables comparisons between sequences and annotations at the genome and base pair level (Carver et al. 2005). Other dedicated synteny views such as SimpleSynteny and Cinteny are capable of visualizing synteny across multiple genomes. SimpleSynteny is a web-based tool providing a pipeline that enables customization of contig organization instead of pure computational predictions for visualizing synteny (Veltri, Wight, and Crouch 2016).

Tools to analyze structural variants (SVs) have also been developed in the past. Ribbon, for example, is a visualization tool developed to support long read evidence in the analysis of structural variation (Nattestad et al. 2020). By displaying long read and whole genome alignments, Ribbon is able to display genomic links that could span several genes going through multiple variants. The general-purpose circular visualization tool Circos (Krzywinski et al. 2009) also supports views that visualize large-scale variation. Copy number variant (CNV) viewers such as the CNSpector can visualize copy number variation and large-scale structural variation that enable the analysis of CNV to detect abnormalities or sequence variants between multiple samples (Markham et al. 2019). General purpose genome browsers such as IGV also remain popular for analyzing SVs (Robinson et al. 2017). An overview of the various visual paradigms of structural variation can be found in (Yokoyama and Kasahara 2020).

## Results

### Advances in JBrowse 2

JBrowse 2 combines the well-established paradigms of general-purpose genome browsers with specialized views of synteny and structural variation. It still uses the fundamental concept of a linear coordinate scheme based on a reference genome, but it also introduces alternative views including circular views, dotplot views, comparative synteny views, and the ability to show discontinuous regions in the linear genome view. This provides a number of different views on structurally disrupted genomes.

Compared to other tools for visualizing structural variants and synteny, JBrowse 2 is most similar to general-purpose genome browsers (GBrowse-Syn and Artemis) in that it renders syntenic relationships between generic linear visualizations of genomes, their annotations, and supporting evidence. However, JBrowse 2 also includes views that draw extensively on user interface concepts pioneered by Ribbon and Circos, and indeed includes many of the views described in (Yokoyama and Kasahara 2020). These are all tied together by a modern web application framework that enables researchers to navigate between these different views, combining multiple coarse-grained, fine-grained, and non-linear views of the genome. In this way, users can begin by visualizing large-scale variation, zoom in to examine a particular feature in detail, and interactively examine the supporting evidence, for example, tracing the local context of a genomic breakpoint.

JBrowse 2 also includes other new features such as the ability to export tracks as publication-quality SVG files; sorting, filtering, and coloring options for alignments tracks; multi-threaded rendering to accelerate the display of multiple tracks at once; and session management, so that users can easily save, restore, export, and share the state of their browser session.

### The JBrowse 2 product range

JBrowse 2 is a family of several apps and modular components produced from the same codebase, specialized for different types of users. These various products are listed in Table 1.

**Table 1.** JBrowse 2 consists of multiple products, aimed at different applications but sharing a common code base.

| Product name | Where to find it | Brief description |
| --- | --- | --- |
| JBrowse Web | https://jbrowse.org/jb2/download/ | Static-site compatible app which can display multiple view types in the same session |
| JBrowse Desktop | https://jbrowse.org/jb2/download/ | Cross-platform desktop app with ability to save user sessions to disk, and display multiple view types in the same session |
| JBrowse CLI | @jbrowse/cli on NPM ("Npm" n.d.) | A command line tool used for administering JBrowse Web instances |
| JBrowse Image CLI | @jbrowse/img on NPM | A command line tool for generating static images (SVG, PNG) of JBrowse sessions |
| JBrowse Embedded Components | @jbrowse/react-linear-genome-view, @jbrowse/react-circular-genome-view on NPM | Libraries that web developers can use to display JBrowse views on their website |
| JBrowseR (Hershberg et al. 2021) | JBrowseR on CRAN (Hornik 2012) | An R package using JBrowse Embedded components that can be used in the RStudio IDE or Shiny apps |
| JBrowse Jupyter (De Jesus Martinez et al. 2022) | jbrowse-jupyter on PyPI ("PyPI · the Python Package Index" n.d.) | A Python package for JBrowse Embedded components that can be used in Jupyter Notebooks |

The two most significant products are the web-based and desktop versions of JBrowse 2, known as “JBrowse Web” and “JBrowse Desktop”. The former runs on any modern web browser; the latter is compiled using the Electron framework to run on macOS, Windows, and Linux. JBrowse Desktop generally has more access to the local filesystem; user’s files can be opened as tracks and will persist across sessions. JBrowse Desktop also works without an internet connection or behind a firewall.

While these apps are mostly identical in look and feel, several key operational details involving sharing and data access are different depending on whether web or desktop is being used. Unless otherwise noted, references to JBrowse 2 in this paper are inclusive of both JBrowse Web and JBrowse Desktop.

### Sessions, assemblies, views, and tracks

Some of the concepts used by JBrowse 2 to organize and integrate different visualizations include *sessions, assemblies, views*, and *tracks*.

#### Sessions

JBrowse 2 uses the term session to represent the current state of the browser. These sessions encompass the state of all views, including the user’s current location in the genome and any data they may have imported. Sessions can be saved, restored, exported, or shared with other users.

#### Assemblies

An assembly in JBrowse 2 refers to a sequence resource, e.g., a FASTA file, and optionally includes a list of aliases describing chromosome names that are to be treated identically, e.g., chr1 and 1. Assemblies can also contain cytoband information that are used to draw ideogram overviews. Multiple assemblies can be loaded at the same time in JBrowse Web and JBrowse Desktop, so a user can load the genome assemblies of multiple species that they want to compare, or different versions of a genome assembly of a single species.

#### Views

JBrowse 2 views are panels that can show data visualizations or other generic things like tabular lists. In JBrowse Web and JBrowse Desktop, the user interface is a vertical arrangement of view panels. By use of these views, different datasets can be arranged next to each other to compare different sets of data, or different visualizations of the same data.

A variety of different views are included with JBrowse 2 in order to accomplish this goal, including the traditional Linear Genome View (Figure 1A & 1B), Circular View (Figure 1C), Dotplot View (Figure 1D), Tabular View (Figure 1E), Linear Synteny View (Figure 1F), and other composite views (Figure 1G & 1H).

**Figure 1:**
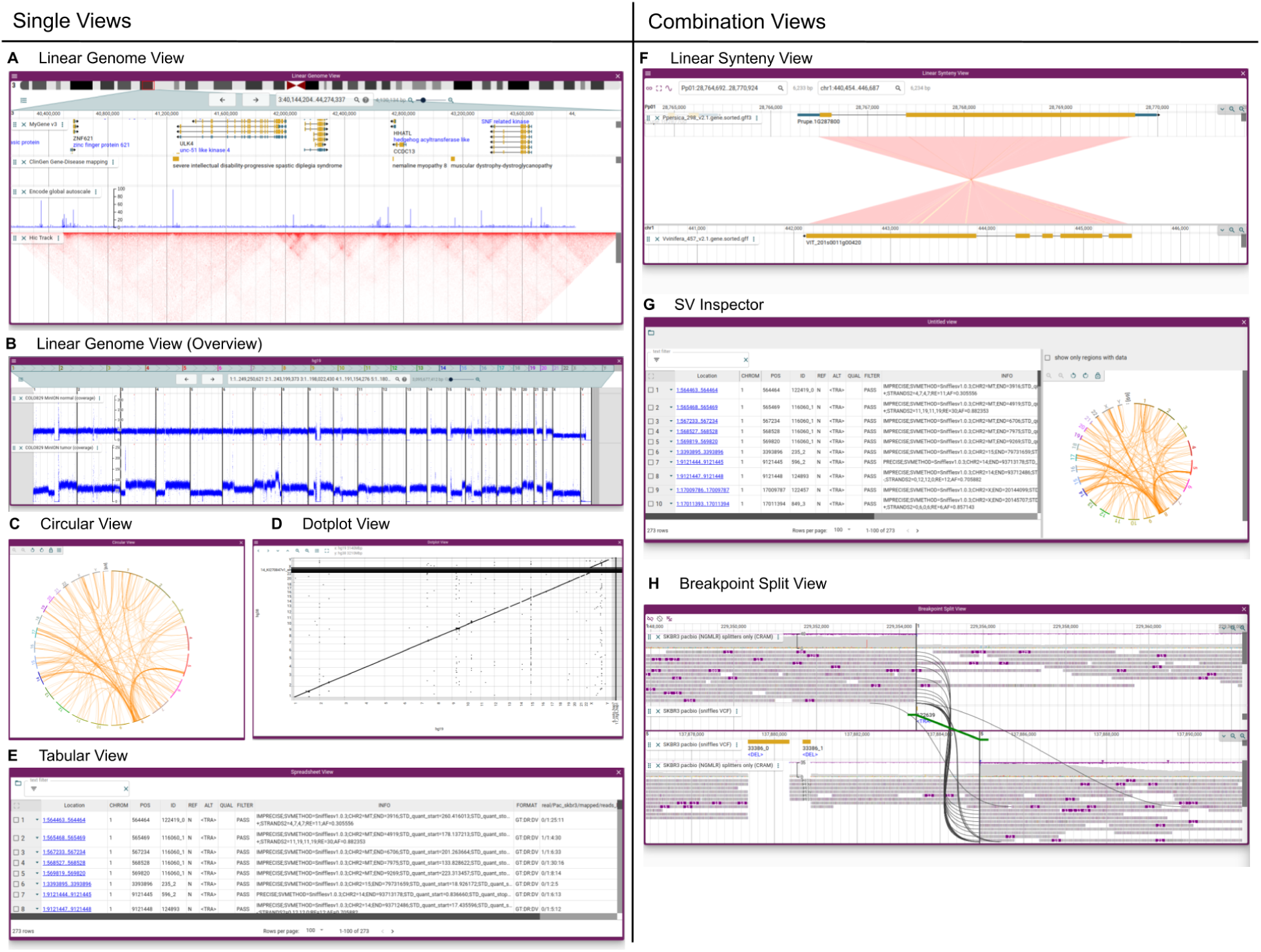
JBrowse 2 integrates many views into a single application. (A) The Linear Genome View displaying gene annotations, quantitative signals, and a Hi-C track. (B) The Linear Genome View can provide a whole genome overview, here showing tumor vs normal sequencing coverage in the COLO829 cell line (Espejo Valle-Inclan et al. 2022). (C) The Circular View gives an overview of long-range relationships within and between chromosomes (here, they are translocations in an SKBR3 cancer genome). (D) The Dotplot View shows relationships between two sequences (in this case the relationship between hg19 and hg38 human genomes). (E) The Tabular View summarizes features in a sortable, filterable list, showing in this example the SKBR3 variant calls from Sniffles. (F) The Linear Synteny View shows relationships between two genomes (in this case, peach and grape) each of which is rendered using a Linear Genome View. (G) The SV Inspector allows inspection of structural variants by combining a Tabular View and a Circular View; here, both the Tabular and Circular views are visualizing a VCF file of translocations in an SKBR3 cancer genome called using Sniffles (Sedlazeck et al. 2018). (H) The Breakpoint Split View shows events such as gene fusions and translocations (in this case, in the SKBR3 cancer genome) by aligning two Linear Genome Views and tracing the split or paired read mappings across the two views.

#### Tracks

Many JBrowse 2 views can display different genome annotation “tracks”: datasets that align in the view and can be selectively hidden, exposed, or reordered by the user. Such annotation tracks are among the earliest established user interface elements in genome browser design, implicitly present in ACeDB (Durbin and Thierry-Mieg 1994) and well-established by the time of GBrowse (Stein et al. 2002) and the UCSC Browser (Kent et al. 2002). A list of track types that are available are listed in Table 2. Tracks can be toggled using the track selector widget.

**Table 2.** The list of available track types in JBrowse 2, which are specialized to render different kinds of data from various sources or file formats. Some of the tracks can be used in multiple view types as well.

| Track Type | Appears in | Function | Supported File Types |
| --- | --- | --- | --- |
| Quantitative Track | Linear Genome View | Displays dense, continuous, quantitative data | BigWig, GC content (from sequence files), GWAS scores (from BED files) |
| Synteny Track | Dotplot View, Linear Synteny View | Displays alignments between different genome assemblies | PAF (Li 2018), .delta from MUMmer (Kurtz et al. 2004), mashmap.out files (Jain et al. 2018), .chain (UCSC), MCScan .anchors files (Tang et al. 2008) |
| Alignments Track | Linear Genome View | Displays a combination of a pileup and a coverage visualization of alignments | BAM, CRAM |
| Hi-C Track | Linear Genome View | Displays Hi-C contact matrix | .hic files, generated by Juicebox (Durand et al. 2016) |
| Variant Track | Linear Genome View, Circular View | Displays feature glyphs corresponding to variants; specialized feature details panel show all genotypes in multi-sample VCF | VCF (plaintext or tabix) |
| Feature Track | Linear Genome View | Displays feature glyphs corresponding to genome annotations, e.g genes | GTF (plaintext), GFF3 (tabix or plaintext), BigBed, BED (tabix or plaintext), features from REST APIs, etc. |
| Reference Sequence Track | Linear Genome View | Displays a reference/assembly sequence and a three-frame translation | FASTA (indexed FASTA or bgzipped indexed FASTA), TwoBit (.2bit) |

### The linear genome view

The Linear Genome View is the primary view in JBrowse 2, and the most similar in look and feel to JBrowse 1, GBrowse and the UCSC Genome Browser. This view shows genome annotation tracks and other genome-mapped data in a horizontally scrollable panel. A screenshot of the Linear Genome View, annotated with key user interface elements, is shown in Figure 2.

**Figure 2:**
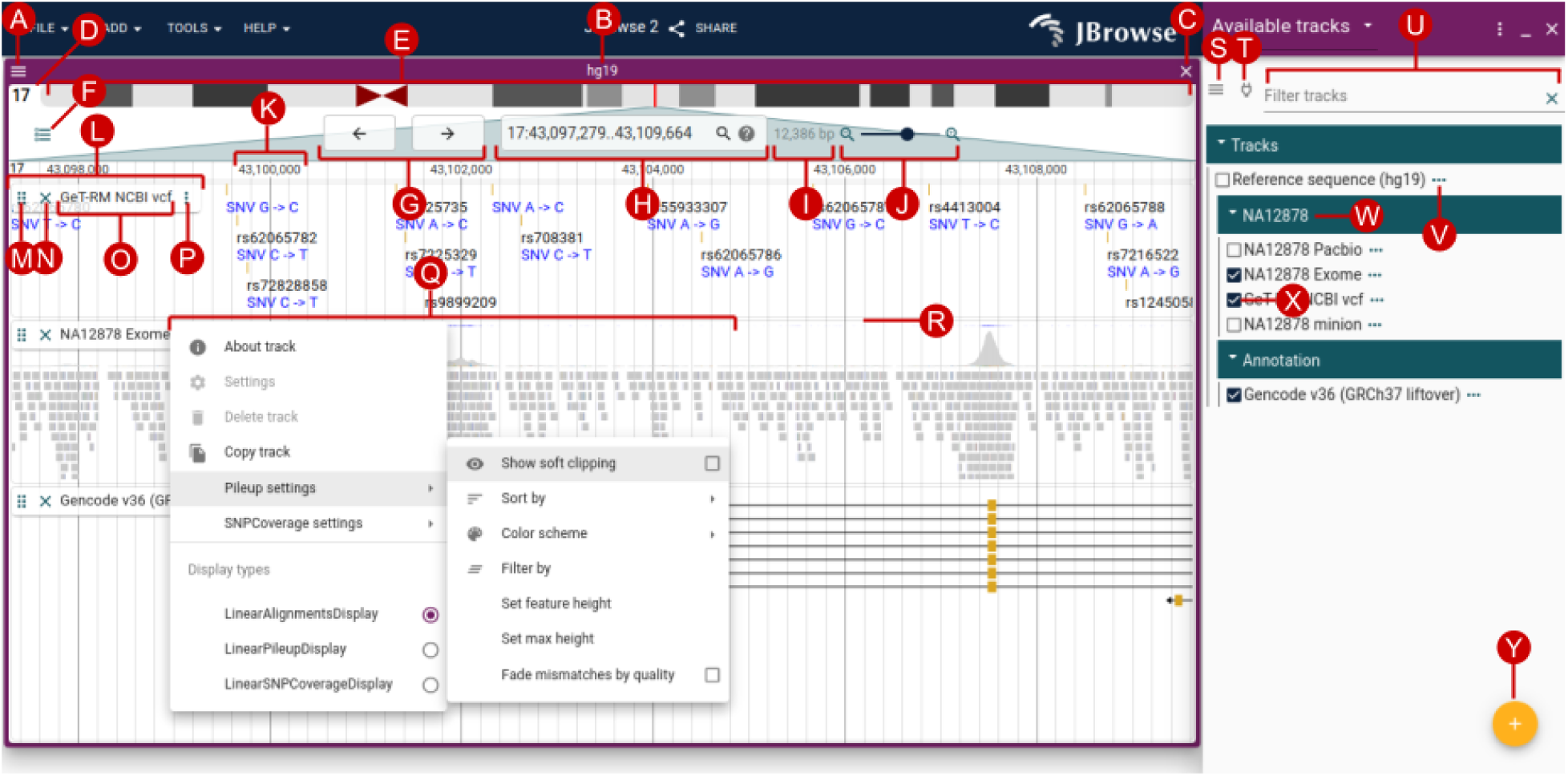
The Linear Genome View is the core view of JBrowse, allowing flexible and interactive examination of a genome sequence and its annotations. The user interface elements annotated on this diagram include (A) view menu, (B) view name, (C) close view button, (D) reference sequence name, (E) reference sequence overview with optional ideogram, (F) open track selector button, (G) pan buttons, (H) location and search box, (I) view size, (J) zoom buttons and slider, (K) major ruler coordinates, (L) track label, (M) track drag handle, (N) track close button, (O) track name, (P) track menu button, (Q) track menu, (R) track resize handle, (S) track selector menu button, (T) connection menu button, (U) track selector filter, (V) track configuration menu button, (W) collapsible category label, (X) track select box, (Y) add track/connection button.

At the top of the Linear Genome View is the navigation bar (Figure 2E–J). Key elements of this area are the currently selected reference sequence, shown either as a ruler or as an ideogram (Figure 2E); navigational controls for panning (Figure 2G) and zooming (Figure 2J); and a location display that doubles as a text search box (Figure 2H).

Beneath the navigation bar is the area where annotation tracks are shown (Figure 2K–R). This area has ruled vertical lines to help see where features are aligned. Handles on the track allow them to be vertically resized, reordered by drag-and-drop, or closed; a track menu exposes more display options and track metadata.

### The track selector

The track selector (Figure 2S–Y) can be used to add or remove new tracks to the current view using a check box. The track selector is associated with one particular view at any given time, so if there are two Linear Genome Views open (e.g., one for the grape genome and one for peach genome), the tracks they display can be configured independently. Furthermore, track selectors are not solely associated with Linear Genome Views; they can also be associated with some of the other views described in later sections, such as the Dotplot View and the Linear Synteny View. For each associated view, the track selector will display tracks relevant to that particular type of view; for example, the track selector for the Dotplot View will only display tracks that are relevant to dotplots.

### Beyond the linear genome view

Complementing the Linear Genome View, several alternate views show different kinds of annotated data, including inter-sequence relationships and large-scale variation. JBrowse 2 provides some mechanisms that link these different views together to facilitate navigation between them; first, through generic inter-view navigation menus (automatically constructed to link alternate views compatible with the same kind of data), and second, through specifically tailored user interface features. For example, right-clicking in an alignments track opens a menu that can launch a Dotplot or Linear Synteny View, as in Figure 7; clicking and dragging a region in a Dotplot View will launch a Linear Synteny View; clicking on breakpoints in the SV inspector will launch the Breakpoint Split View, and so on.

### Displaying and comparing multiple assemblies

JBrowse 2 features several specialized synteny views, including the Dotplot View and the Linear Synteny View. These views can display data from Synteny Tracks, which themselves can load data from formats including MUMmer (Kurtz et al. 2004), minimap2 (Li 2018), MashMap (Jain et al. 2018), UCSC chain files (“Chain Format” n.d.), and MCScan (Tang et al. 2008).

The Dotplot View (Figure 3) can be used at different zoom scales to display whole-genome overviews of synteny, close-ups of individual syntenic regions, and even individual long reads aligned to the reference sequence (see “Visualizing long reads”). Users can click and drag on the Dotplot View to open a Linear Synteny View of the region.

**Figure 3:**
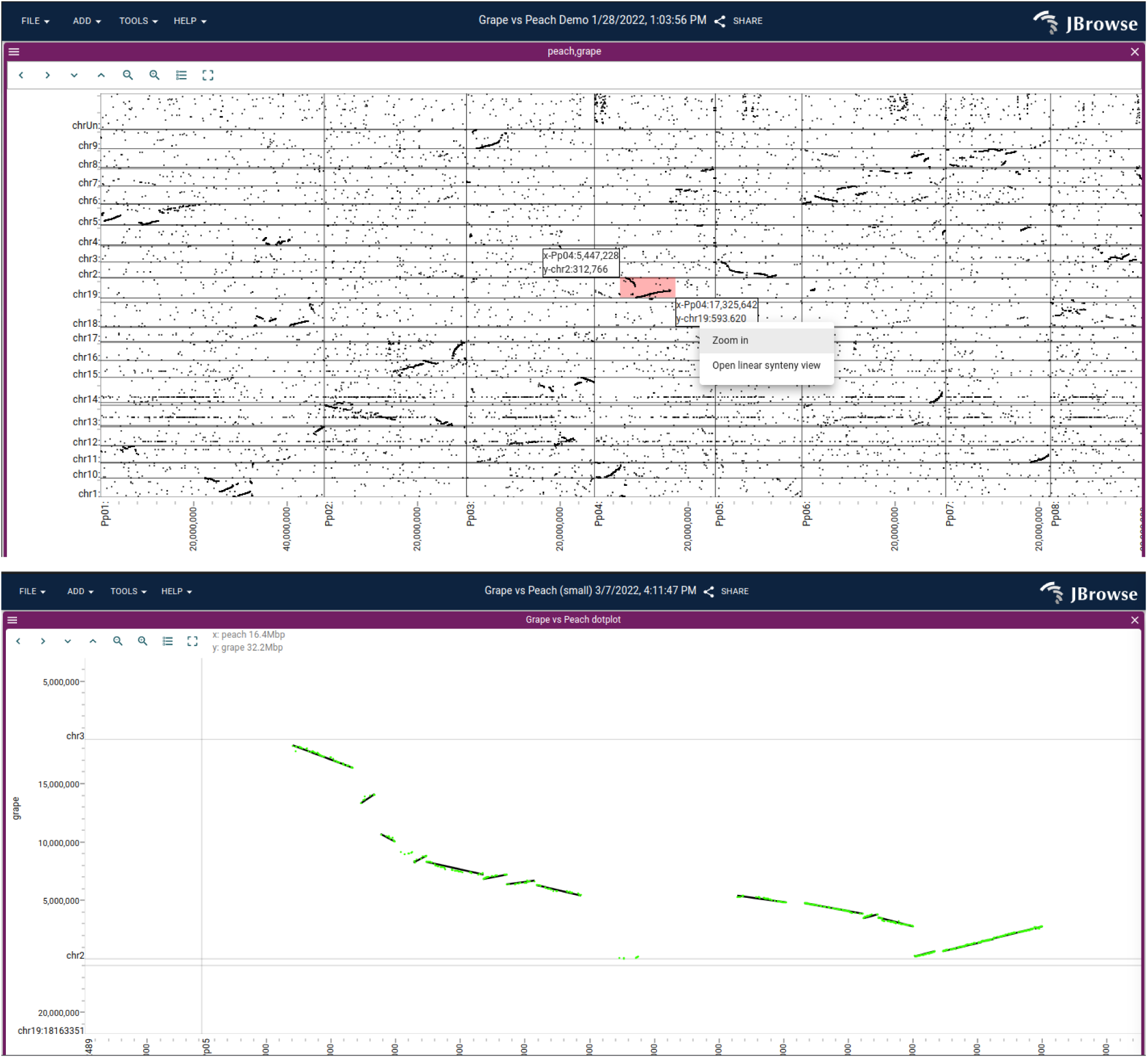
(A) A dotplot showing a whole-genome alignment of the grape vs peach genome, computed by minimap2 and loaded in PAF format, reveals large-scale syntenic structure. The user can click and drag on this view, highlighting an area shown by the small pink rectangle, to open up a detailed view (B) showing multiple synteny tracks, with individual gene pairs (green) and larger syntenic blocks (black) from MCScan.

The Linear Synteny View (Figure 4A) shows two linear genome view panels stacked vertically. This feature allows users to view Synteny Tracks, representing regions of similarity between two different assemblies. The top and bottom panels are each fully featured Linear Genome Views, to which annotation tracks can be independently added. In addition, by exploiting the feature of the Linear Genome View whereby discontiguous regions can be shown, the user can view distal gene duplications within the Linear Synteny View (Figure 4B).

**Figure 4:**
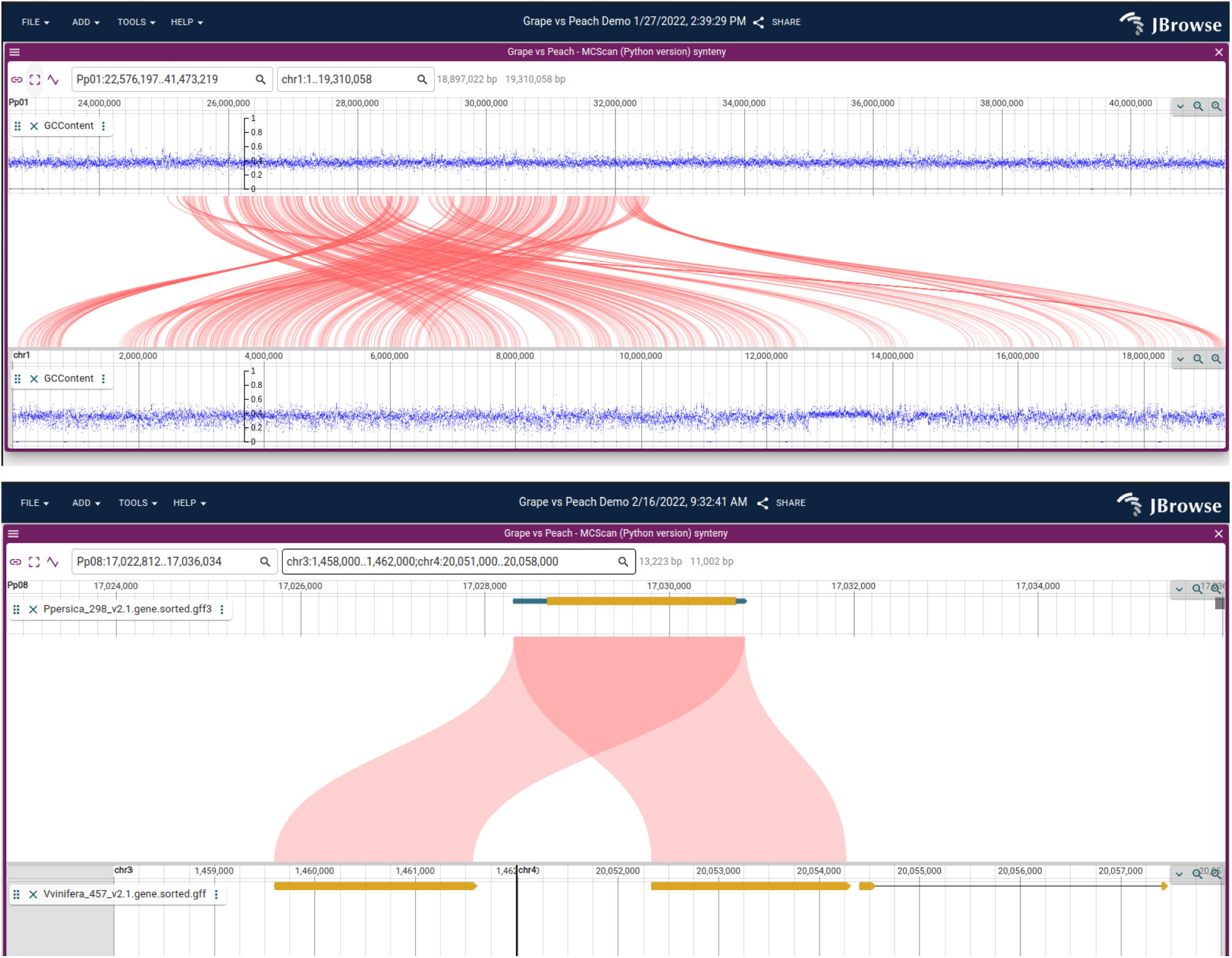
(A) The Linear Synteny View comparing the grape and peach genomes using data from MCScan reveals a complex rearrangement. (B) A close up view of a gene duplication visualized with the Linear Synteny View, with discontinuous regions (chr3 and chr4) displayed side by side on the bottom panel.

### Displaying structural variation

Structural variants can be classified into simple types (e.g., duplications, inversions) and more complex types arising from combinations of the simpler ones. The visualization of such SVs is challenging because the derived genome (e.g., the genome incorporating the structural variant) may be significantly different from the reference genome. As a result, it is often appropriate to use different visualization modalities depending on the type of SV.

#### Overviews of structural variation

The availability of multiple view types can help users visualize SVs through different lenses. For example, whole-genome overviews are often helpful to visualize large-scale patterns of structural variation. The Linear Genome View can be used to get a quick visualization of copy number variation by visualizing read depth from BigWig files representing genome sequencing coverage, employing its facility to display multiple chromosomes side-by-side to get a whole-genome overview (Figure 1B).

Users can also apply one of the specialized JBrowse 2 views designed for whole-genome or multi-genome overviews. These include the Circular, Tabular, and SV inspector views.

The Circular View displays annotations in a circular format as popularized by Circos (Krzywinski et al. 2009). Because of its compact arrangement, this circular view is beneficial for exploring long-range structural variations encoded as breakends or translocations in VCF files (“The Variant Call Format Specification - VCFv4.3 and BCFv2.2” 2022), BEDPE files, or STAR-fusion (Haas et al. 2017) results.

The Tabular View is different from other views described in that it is a textual list of features rather than a graphical visualization. The table columns show key fields from the variant file, such as the type of SV, the location, and the ID. Controls in each column allow the tabular view to be filtered or sorted to drill down into the variant list.

The SV Inspector (Figure 5) combines the Circular View and the Tabular View to allow users to prioritize their structural variants. This composite view was developed to address a common workflow in cancer bioinformatics research: examining a list of putative variants to visually evaluate them in the context of relevant information such as canonical gene models, RNA-seq results, chromatin interactions, or other genome annotation data. The SV Inspector includes a Tabular View of a set of candidate structural variants with controls to mark features for later inspection and a Circular View visualizing where these variants lie in the genome. Each variant is presented as a row in the tabular view and as a chord in the circular view. Clicking on a chord in the Circular View or a row in the Tabular View launches a Breakpoint Split View (Figure 6) showing the read evidence for a selected structural variant.

**Figure 5:**
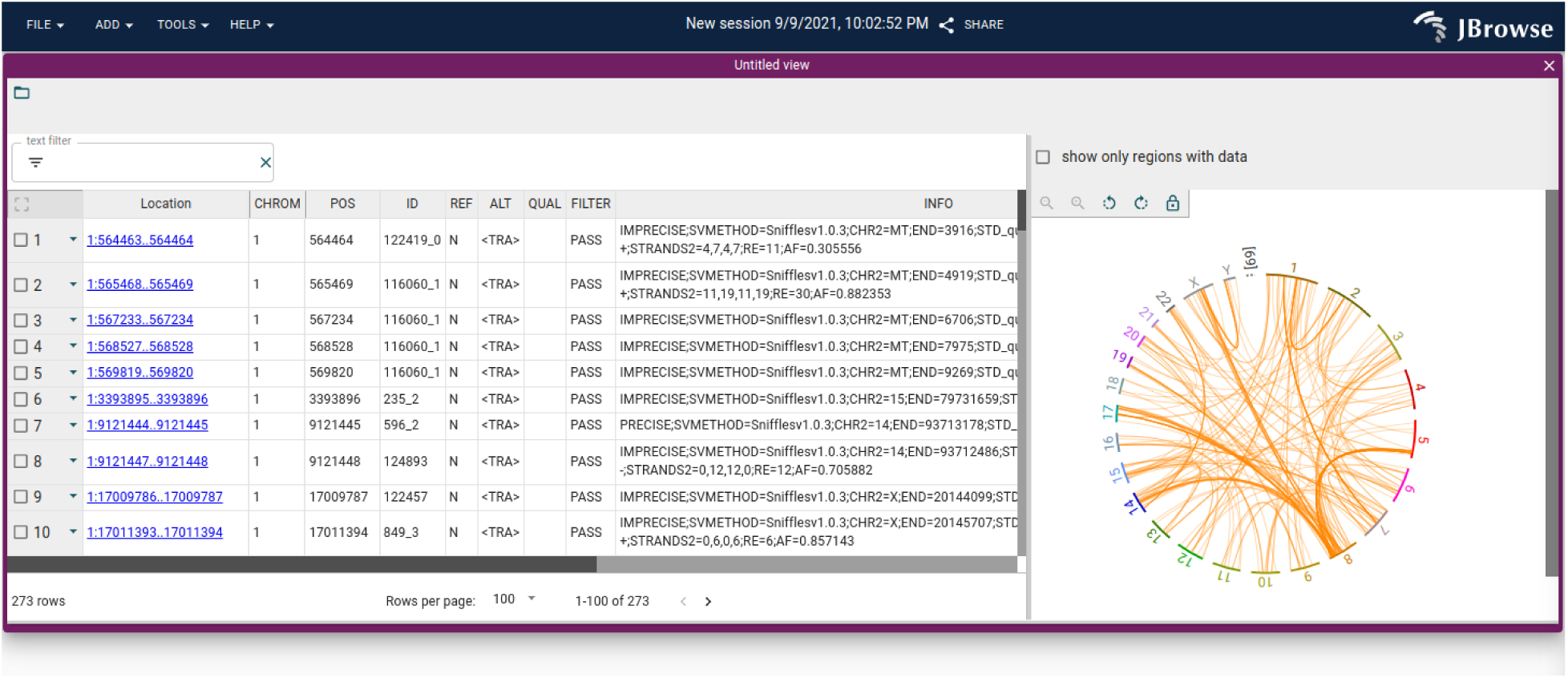
The SV Inspector showing structural variants in the SKBR3 breast cancer long read dataset. The SV Inspector places the Circular and Tabular views side-by-side. On the left, the Tabular view can be filtered using simple text expression filters (text box at top left), or column filters (controls in each column header). The results of filtering are reflected in the circular view at right.

**Figure 6:**
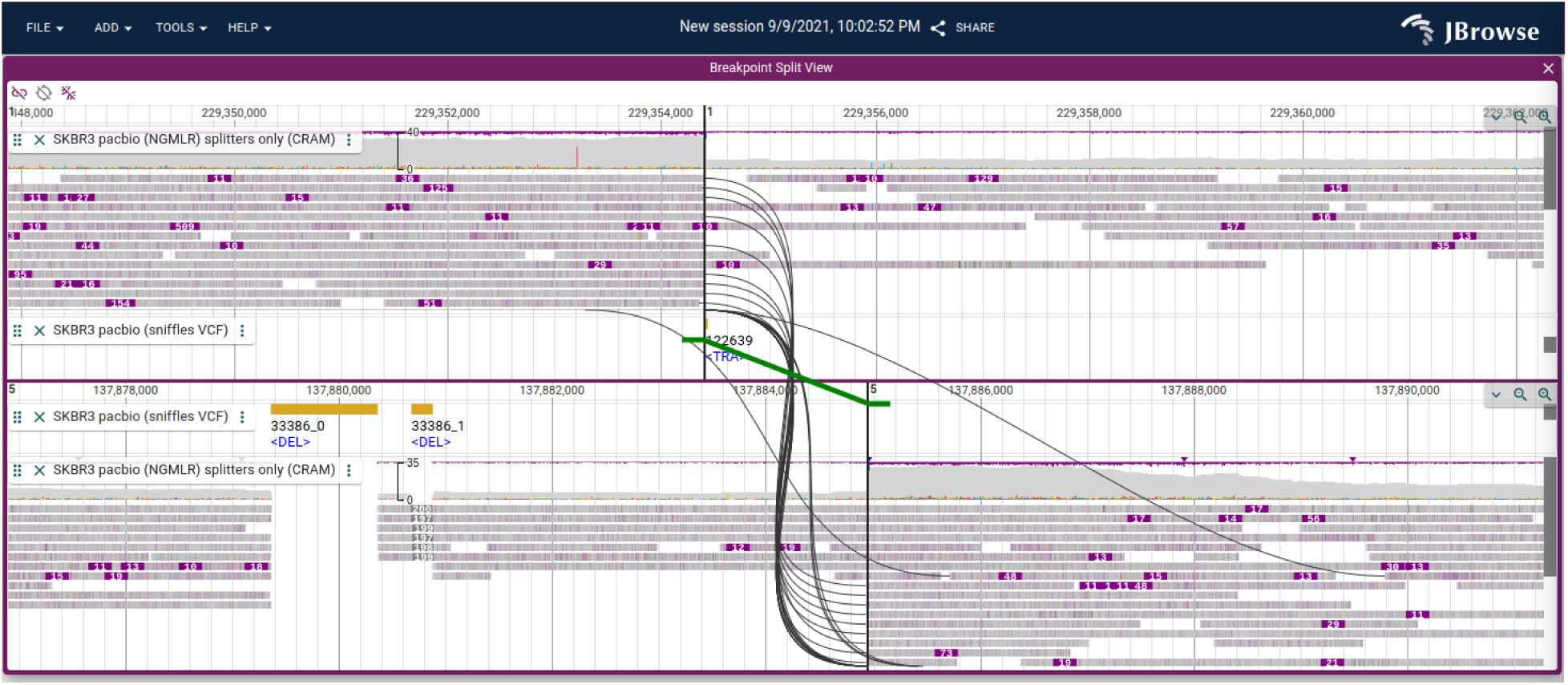
A Breakpoint Split View showing a chromosomal translocation connecting chr1 and chr5 in the SKBR3 breast cancer cell line. The split long read alignments are connected using curved black lines, and the variant call itself is shown with a green line with directional “feet” showing which sides of the breakpoint are joined.

#### Fine detail of structural variation

To visualize a single breakpoint, such as a gene fusion and the evidence for that breakpoint, we introduce the Breakpoint Split View. The Breakpoint Split View consists of two Linear Genome Views stacked vertically. The power of this view lies in its ability to visualize genomic evidence for structural variants between discontinuous regions of the genome (Figure 6). The read evidence for the structural variant is shown using curved black lines for long split alignments or paired-end reads. The structural variant call itself is shown using green lines, based on information from breakends (“The Variant Call Format Specification - VCFv4.3 and BCFv2.2” 2022) or translocation type features from VCF files.

### Visualizing long reads

Long read sequencing technology, such as the platforms developed by Pacific Biosciences and Oxford Nanopore Technologies, has proven useful in the resolution of haplotypes, structural variants, and complex repetitive regions. JBrowse 2 includes several features to highlight the information contained in long reads, including: advanced alignments track features to sort, filter, and color reads and a “read vs reference” feature that enables users to view alignments of long reads in a Dotplot View or Linear Synteny View (Figure 7).

**Figure 7:**
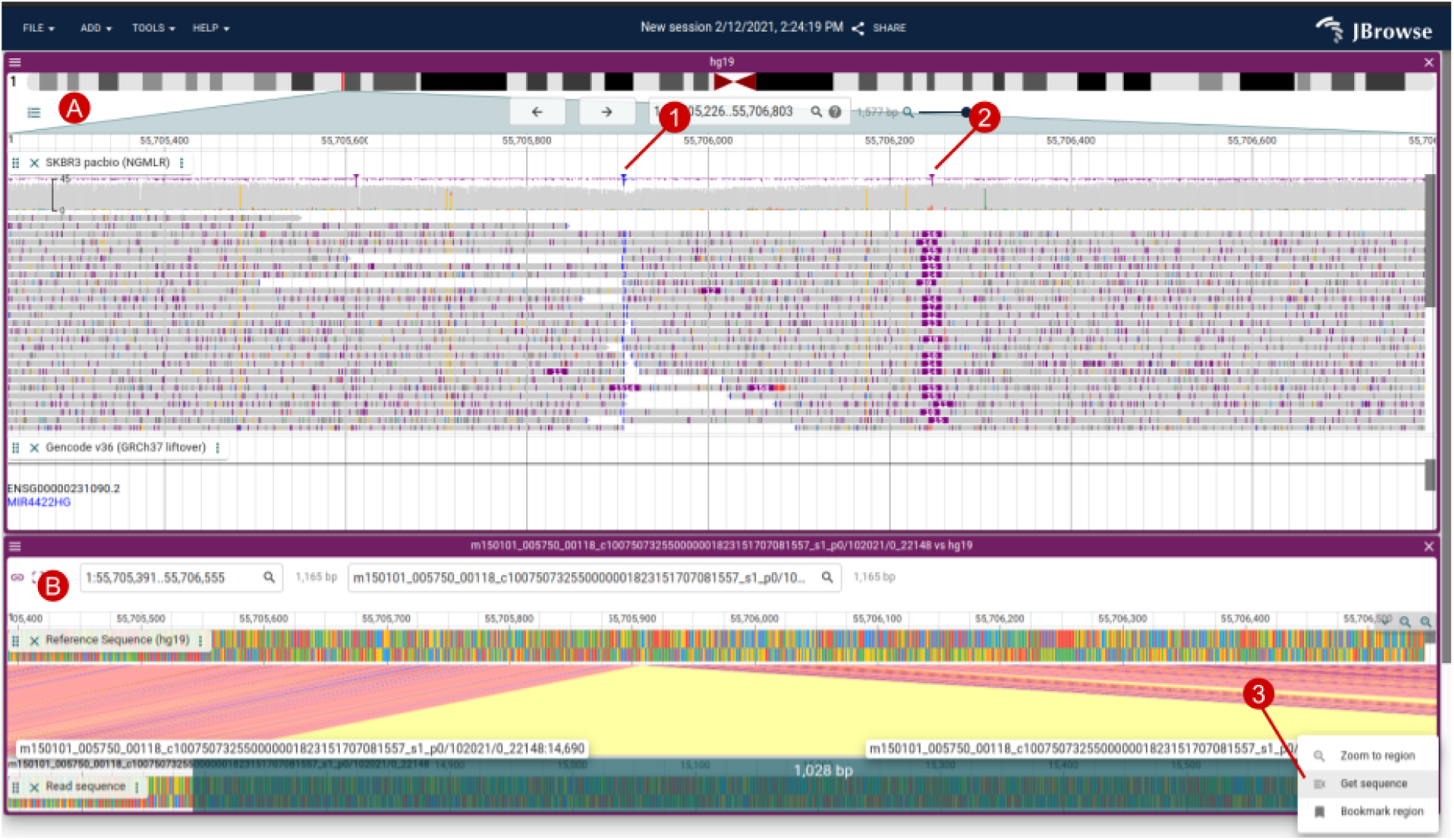
(A) An alignments track showing SKBR3 pacbio alignments, with a large (>1000bp) insertion highlighted by soft-clipped reads in blue (1) and a smaller insertion in purple (2) (B) The “Read vs ref” view created by right clicking a read in the Alignments Track creates a Linear Synteny View comparing the read vs the reference genome, which allows you to see the inserted non-reference bases easily. Users can also select regions on the read or reference sequence and select “Get sequence” (3).

In addition to elucidating long-range structure, long-read sequencing platforms can provide a direct readout of chemical modifications such as methylation on DNA and RNA sequences. The modifications can be called by tools such as nanopolish (Loman, Quick, and Simpson 2015) and primrose (*Primrose: Predict 5mC in PacBio HiFi Reads* n.d.), which stores this information in the MM tag in BAM/CRAM files (“HTS Format Specifications” n.d.). JBrowse 2 can then use the MM tag to render the positions of these modifications on individual reads.

### Ways to access data

#### Connections

JBrowse 2 allows users to create “Connections” to sets of tracks or assemblies that are understood to be managed outside of the JBrowse instance. JBrowse 2 includes two types of Connections by default: JBrowse 1 Connections, which allow viewing tracks from a JBrowse 1 installation on the web, and UCSC Track Hub Connections, which allow viewing tracks in a UCSC Track Hub (Raney et al. 2013). To help users navigate the latter type of Connection, JBrowse 2 includes an interface for browsing the UCSC Track Hub Registry (The Ensembl Core Team n.d.).

#### Direct access to local files

JBrowse Web and JBrowse Desktop allow users to open tracks directly from a user’s local filesystem. This functionality keeps the data private on the user’s computer. On JBrowse Web, due to limitations of web browsers, local files must be re-opened when the page refreshes or a session is re-opened. JBrowse Web provides a message to alert users to this necessity. This limitation does not exist on JBrowse Desktop.

#### Sharing sessions

In JBrowse Web, users can share sessions with other users by generating a share link, which produces a shortened URL containing the contents of the users session. Visiting the link will restore all the same views at the same locations, with the same tracks displayed. This also includes track data that a user has added to the session: if a user opens a track in their session that references a remote file, then their share link will include this track.

#### Authentication

JBrowse 2 natively supports Google Drive, Dropbox, and HTTP Basic authentication. Plugin developers can extend this to connect the genome browser to any application that needs authentication to access data, as described in the section titled “Extending JBrowse 2 with Plugins”.

### Performance and scalability

JBrowse 2 is structured to take heavy computations off the main thread using remote procedure calls (RPC). In JBrowse Web and JBrowse Desktop, we use web workers to handle RPCs, which perform data parsing, rendering, and other computationally time-consuming tasks in a separate thread. This approach allows the app to remain responsive to the user even when displaying large datasets or multiple tracks. Note that JBrowse Embedded does not use web workers currently, so it is single threaded, but may gain web worker support in the future.

To profile the end-to-end performance of loading and rendering tracks, we used Puppeteer (*Puppeteer: Headless Chrome Node.js API* n.d.) to run JBrowse 2 (both with parallel rendering enabled and disabled), JBrowse 1, and igv.js. Each tested browser was given the task of rendering BAM and CRAM files containing long and short reads at varying coverage. Full details of the benchmark can be found under “Performance and Scalability benchmark details” in the Methods section.

When rendering a single track, JBrowse 2 has comparable performance to igv.js (Robinson et al. 2020), as shown in Figure 8. However, when rendering multiple tracks, the parallel rendering strategy in JBrowse 2 can improve performance (Figure 9). The parallel strategy also yields a more responsive user interface, since the main thread (whose frame rate determines the apparent speed with which the browser responds to user input) does not become tied up by rendering and data parsing (Figure 10).

**Figure 8:**
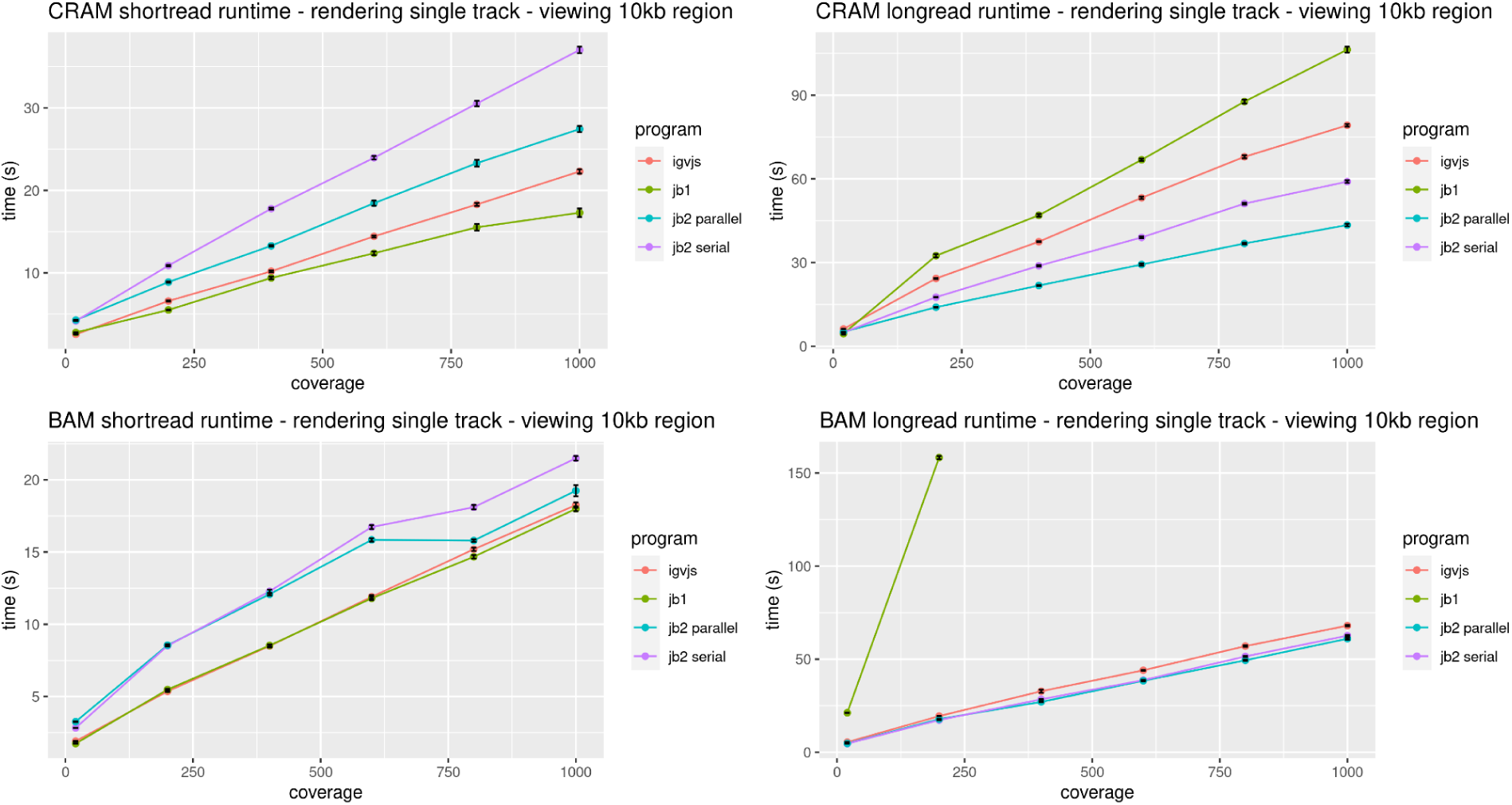
JBrowse 2’s performance is comparable to igv.js, and significantly exceeds JBrowse 1’s performance on large datasets, with modest improvements when running in parallel, as reflected in these benchmarks rendering aligned reads of varying coverage and file formats in a 10kb region. The incomplete data for JBrowse 1 on the BAM long read benchmark reflects the fact that JBrowse 1 times out this benchmark on dense coverage datasets (i.e. its rendering time exceeds 5 minutes). Full details of the benchmark can be found under “Performance and Scalability benchmark details” in the Methods section.

**Figure 9:**
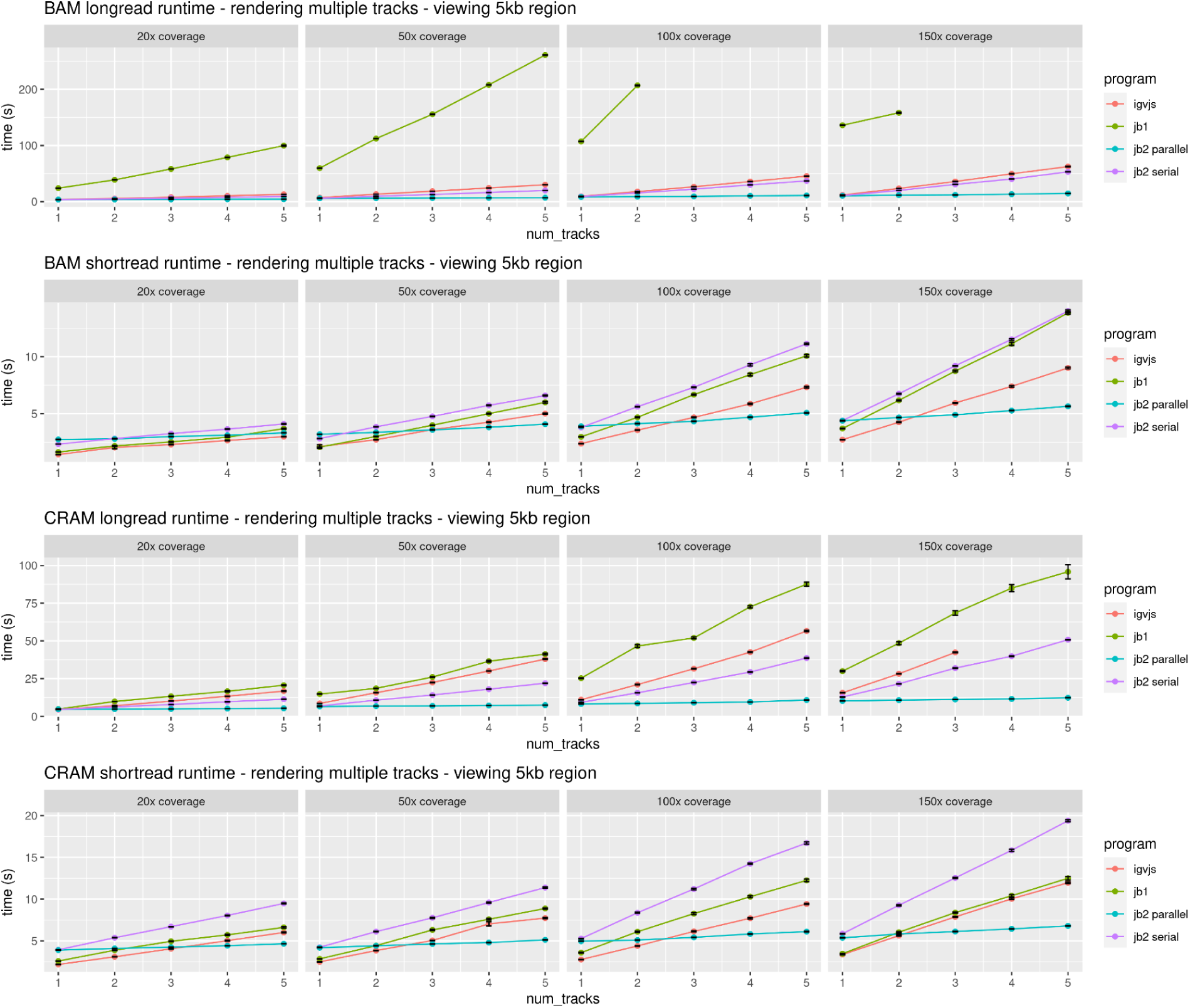
When displaying multiple tracks at once, JBrowse 2’s parallel rendering strategy shows significant gains compared to single-threaded (serial) strategies, as reflected in these benchmarks rendering aligned reads of varying coverage and file formats in a 5kb region. The incomplete data for JBrowse 1 and IGV.js on some of the benchmarks reflects the fact that these apps time out our benchmark under some circumstances (i.e. rendering time exceeds 5 minutes). Full details of the benchmark can be found under “Performance and Scalability benchmark details” in the Methods section.

**Figure 10:**
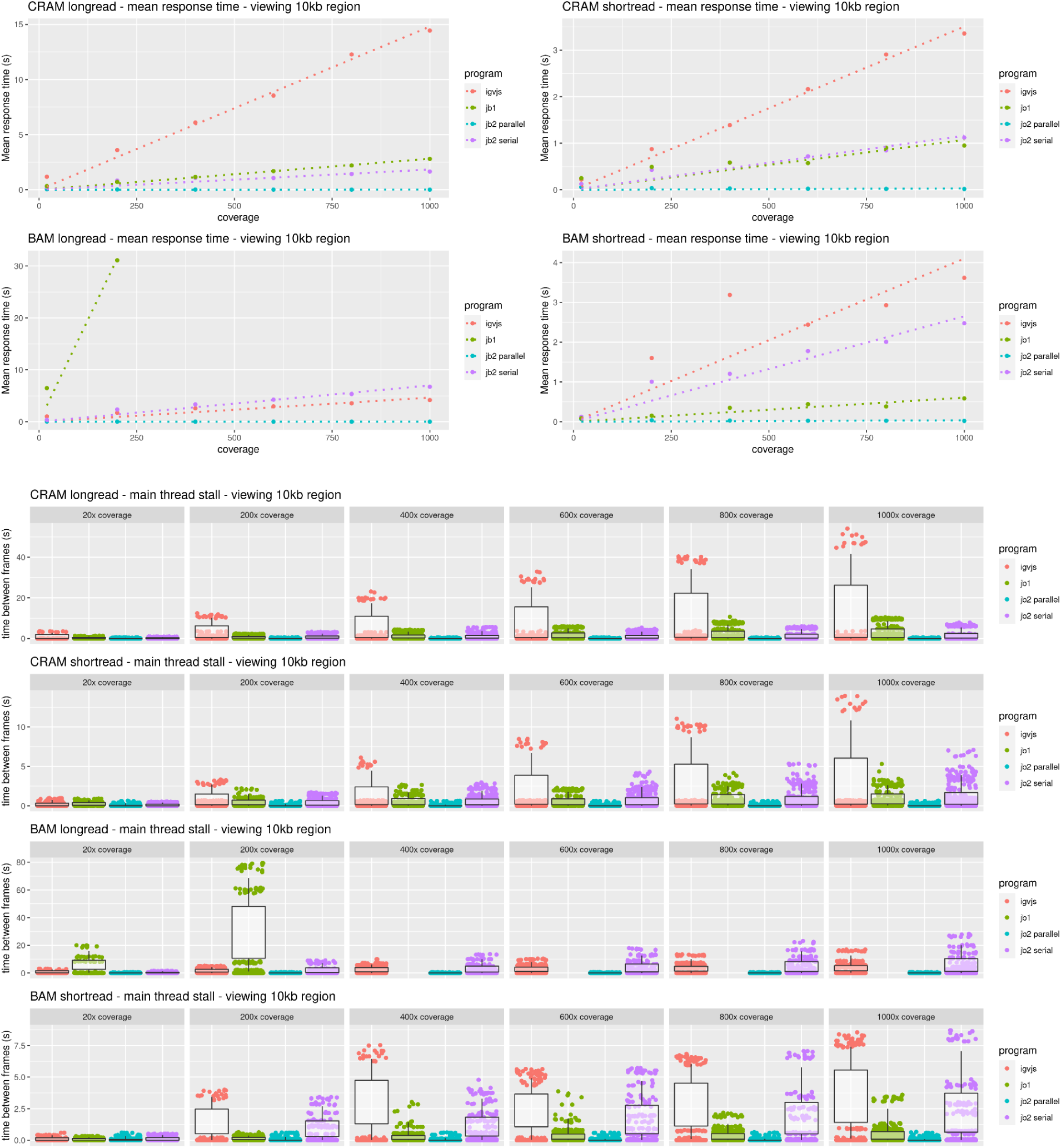
JBrowse 2’s parallel rendering strategy yields significant improvements in user interface responsiveness, as reflected in these benchmarks rendering aligned reads of varying coverage and file types in a 10kb region. We define the *response time* as the delay, during the rendering phase of the benchmark, from a randomly sampled time point until the time the next frame is rendered. This directly reflects the perceived delay between when a user initiates an action and when the app responds. The response time is a random variable; its expectation gives a sense of average lag, while its variation gives a sense of how unpredictable the user interface delays can be. Panel (A) plots the expectation of the response time as a function of sequencing coverage. At high coverage, JBrowse 2’s parallel strategy maintains a low response time, in contrast to single-threaded strategies whose response time can grow large, with the perception that the browser is “hanging” or “frozen”. The relationship between response time and coverage is approximately linear for all browsers, as shown by the dotted linear regression fit. The incomplete data for JBrowse 1 on the BAM long read benchmark reflects the fact that the simulation times out (rendering time >5 minutes). Panel (B) shows the same data plotted as a scatterplot of time between frames. The plotted points show the raw time between frame values, and are overlaid with boxplots that show the variation in response times (25th and 75th percentiles shown in the boxes, 5th and 95th percentiles shown in the tails). Full details of the benchmark can be found under “Performance and Scalability benchmark details” in the Methods section.

### Administration and configuration

#### Extending JBrowse 2 with Plugins

JBrowse 2 has a plugin system which provides developers with the ability to customize and extend JBrowse to suit the specialized needs of the organization or researcher. Table 3 describes the elements that can be extended via plugins, such as data adapters, track types, and view types.

**Table 3.** A listing of JBrowse 2 elements that can be extended by third-party plugins.

| Plugin-Extensible Element | Description | Example |
| --- | --- | --- |
| Data adapters | Classes through which reading and parsing unique data formats is done, including retrieving data from RESTful APIs. | BamAdapter: processes .bam files, remote or local, along with their coordinating .bai or .csi index files |
| Text search adapters | Classes through which searches for features by name are processed. | TrixTextSearchAdapter: used for UCSC trix indexes, generated by the jbrowse text-index command |
| Track types | A high level concept in the configuration system that associates a name, trackId, and metadata with a data adapter. There is not a lot of logic attached to track types; instead, the display types and renderers are used to draw and add logic to tracks. | AlignmentsTrack: displays data typically associated with BAM and CRAM type data adapters |
| Display types | The code to “display” a track type in a particular view. This layer is important because it allows track types to be displayed in multiple view types, and often contains logic such as menus, click actions, React components, and more. | LinearSyntenyDisplay, DotplotDisplay: these display types help display a SyntenyTrack in different view types |
| Renderer types | Fetches data from a data adapter and draws the features, typically rendering to either SVG or HTML5 canvas. Renderers run on the remote side of an RPC call, and are instructed to render a particular genomic region or regions. | PileupRenderer: draws the reads from a .bam or .cram file |
| Widgets | User interfaces that provide utility or information to the user. In JBrowse Web and JBrowse Desktop, these appear as a side drawer. In embedded, these appear in a dialog box. | Track selector: Provides a list of tracks for user to toggle in the user interface |
| RPC calls | Custom code that a plugin runs on the remote side of an RPC call (e.g., in a web worker) to avoid doing heavy work in the main thread. | WiggleGetMultiRegionStats: estimates quantitative statistics for the given genomic regions |
| View types | Containers for visualizations that permit wholly unique visualizations to be displayed in the same context as the standard linear genome view. | CircularView: provides a circular whole-genome overview of structural variants |
| Extension points | A named list of callbacks that the code can use to create arbitrary modifications to various parts of the code. | The ``extendPluggableElement`` extension point allows plugins to add to state tree models e.g., add menu items |
| Connection types | Connections provide a way to connect to a provider of tracks or assemblies. | UCSCTrackHubConnection: adds tracks from a UCSC track hub to the tracklist |
| Internet account type | Adds a custom authentication method to a remote data provider | GoogleDriveOAuthInternetAccount: Adds ability to open share links from Google Drive |

JBrowse Web and JBrowse Desktop feature an in-app Plugin Store where users can install plugins. Examples of third-party plugins that can be installed include a multiple sequence alignment viewer, an ideogram viewer with Reactome (Gillespie et al. 2021) pathway visualization (Figure 11), and plugins that provide data adapters for fetching data from the mygene.info (Xin et al. 2016) and the CIVIC API (Griffith et al. 2017).

**Figure 11:**
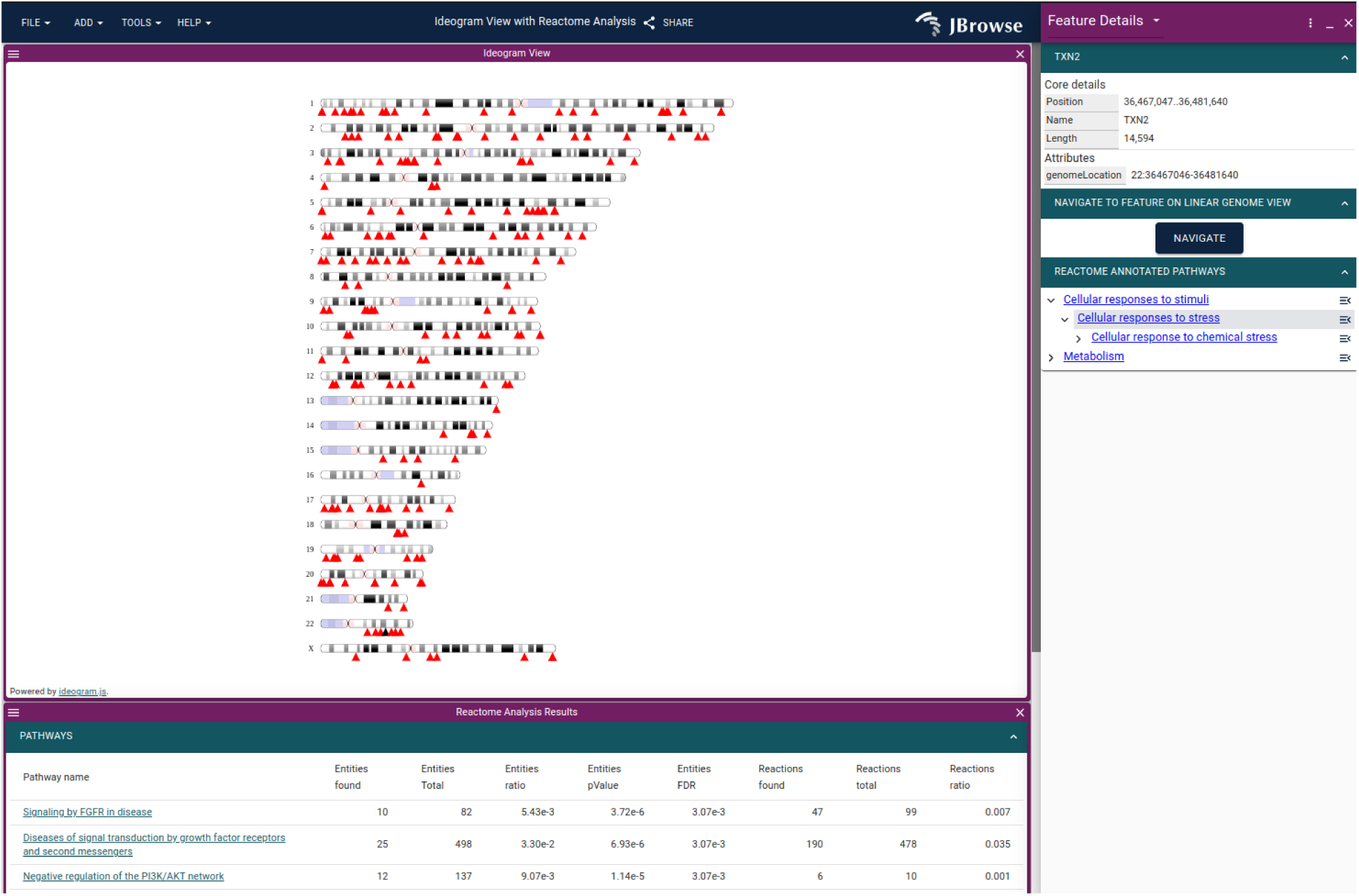
The ideogram view showing the Reactome pathway analysis on a gene list using the JBrowse 2 Ideogram plugin, which uses ideogram.js (Weitz n.d.).

#### Static site compatibility

JBrowse 2 is a “static site” compatible application: since all data parsing and rendering occurs on the client, its ongoing deployment does not require any server-side code beyond a basic web server. Static sites are low-cost because they can be hosted on inexpensive or free hosting services like Amazon S3 and Github pages. In addition, many security issues are mitigated, and the sites may require less maintenance.

#### JBrowse CLI

The JBrowse CLI is an administrative tool that can load assemblies, tracks, and indices for gene name searching. The JBrowse CLI tool can be installed from NPM and runs on both Unix-like and Windows systems, somewhat more portably than JBrowse 1, which required Perl scripts whose installation on Windows-like systems was more involved. The JBrowse CLI also includes an admin-server command which allows changes made in the web GUI to be persisted to the config file on disk.

#### Text indexing

The JBrowse CLI includes a text indexing command that creates trix formatted indices (“Trix Indices” n.d.); these indices are easier to manage than the index files used by JBrowse 1 (which consisted of an on-disk hash table built from many small files). The text indexing in JBrowse 2 allows for either per-track indexes or aggregate indexes containing data from multiple tracks. The tool can index gene IDs, full-text descriptions, or other arbitrary data fields from GFF and VCF files. Text searching can also be extended using plugins to adapt to custom search systems.

## Discussion

In this paper, we have introduced JBrowse 2. This document is not intended to recap the complete JBrowse 2 user guide; instead, it covers foundational concepts (sessions, assemblies, views, tracks, connections, and plugins), the various views (linear genome view, other basic views, composite views, and overviews), and the data access modalities, with a focus on use cases involving comparative genomics and structural variation. We have also briefly introduced some technical details (including administration, performance engineering, and plugin development) and deployment on non-web platforms, including R, Jupyter/Python, and desktop.

In large part, the potential of JBrowse 2 arises from its portability, extensibility and customizability. It can run as a desktop application for personal use, as a web app to support research groups and communities, as a visualization component within Jupyter Notebooks or R for use by data scientists, or as a series of embedded components for developers of biomedical data portals and other integrative projects. In addition, JBrowse 2 is customizable at multiple levels, ranging from an intuitive GUI for view and track configuration, to scriptable customization via its CLI, to bespoke visualizations and user interfaces created using a developer’s plugin API.

## Methods

### Software methodology

The JBrowse development team follows agile practices to plan, design and implement new software. Requests for new features and bug reports are documented using Github issues and reviewed by the development team during backlog grooming sessions. Teammates hold pair programming sessions to discuss and review implementations. Pull requests with new features are peer reviewed by developers to test and approve changes. A Github project board is used to track in progress issues and organize incoming ones according to priority. Finally, a test suite is run for all changes to the codebase using Github Actions. All source code for JBrowse 2 is distributed under the Apache License version 2.0 (“Apache License, Version 2.0” n.d.). Many utility libraries and data parsers that are used by the JBrowse project are also published on NPM, which can be used independently of JBrowse itself.

### Software design sprints

We held two design sprints focused on specific use cases of JBrowse 2, collaborating with members of the existing JBrowse 1 user community and other selected groups. The first took place at the Ontario Institute for Cancer Research in Toronto in 2019, focusing on prioritization of structural variants. The second took place remotely over Zoom in 2020, and focused on comparisons between genomes. Both of these sprints approximately followed the Google Ventures design sprint model. For a week, working in small teams, we conducted expert interviews, mapped out user journeys, sketched out competing solutions, built prototypes, and then presented these prototypes to users for testing.

### Implementation details

JBrowse 2 relies on several technological innovations that were not available when JBrowse 1 was created. JBrowse 2 uses TypeScript (“JavaScript With Syntax For Types” n.d.), which adds compile-time type checking to JavaScript. JBrowse 2 also uses React (“React” n.d.) for rendering the user interface and mobx-state-tree (“Welcome to MobX-State-Tree!” n.d.), which provides a centralized way of storing, accessing and restoring the application state. In addition, JBrowse Web and JBrowse Desktop use Web Workers (“Using Web Workers” n.d.) to enable parallel processing and rendering of data tracks. JBrowse Desktop is built using Electron (“Electron” n.d.), a system for building cross-platform desktop applications.

### Performance and scalability benchmark details

For performance profiling, we generated reads from an arbitrary region chr22:25000000-25250000 on hg19. We simulated 50kb long reads to a coverage of ∼1000x using pbsim2 v2.0.1 (‘pbsim ref.fa --depth 1000 --hmm_model data/R103.model --length-mean 50000 --prefix 1000x’) and also simulated 150bp paired-end short reads to a coverage of ∼1000x using wgsim v1.15.1 (‘wgsim -1 150 -2 150 -N 1000000 hg19mod.fa 1000x.1.fq 1000x.2.fq’). We aligned the reads to the genome using minimap2 (2.24-r1122), and subsampled the resulting different coverages using samtools (1.15.1). We instrumented igv.js (v2.12.1) to output a console log when it completed rendering. We then compared the timing of igv.js with JBrowse 2 Web (v1.7.7), JBrowse 2 Embedded (v1.7.7), and JBrowse 1 (v1.16.11). The benchmarking script uses puppeteer v3.16.0 (*Puppeteer:* Headless Chrome Node.js API n.d.), and measures the time taken to complete the rendering of a track. Each step is run N=10 times, and the mean time with standard error bars (SE = σ/sqrt(N)) are plotted. The frame rate is calculated using the requestAnimationFrame API. A reproducible benchmark script is available at https://github.com/cmdcolin/jb2profile.

## Availability of data and materials

The datasets analyzed during the current study are listed below, with links from which they can be downloaded and citations to the primary scientific publications associated with the data.

- The grape genome (Vvinifera_145_Genoscope.12X.fa) and annotations (Vvinifera_457_v2.1.gene.gff3) are available from Phytozome https://phytozome-next.jgi.doe.gov/info/Vvinifera_v2_1 (Le Clainche Giorgio Malacrida Eléonore Durand Graziano Pesole Valérie Laucou Philippe Chatelet Didier Merdinoglu Massimo Delledonne Mario Pezzotti Alain Lecharny Claude Scarpelli François Artiguenave M. Enrico Pè Giorgio Valle Michele Morgante Michel Caboche Anne-Françoise Adam-Blondon Jean Weissenbach Francis Quétier & Patrick Wincker 2007)
- The peach genome (Ppersica_298_v2.0.fa) and annotations (Ppersica_298_v2.1.gene.gff3) are available from https://phytozome-next.jgi.doe.gov/info/Ppersica_v2_1 (Verde et al. 2013)
- The SKBR3 breast cancer cell line PacBio sequencing and Illumina sequencing are available from http://schatz-lab.org/publications/SKBR3/ (Nattestad et al. 2017)
- The COLO829 melanoma cancer cell line Nanopore sequencing is available from ENA PRJEB27698
- Whole genome alignment of grape vs peach generated with minimap2 2.24-r1122 (‘minimap2 -c Vvinifera_457_Genoscope.12X.fa.gz Ppersica_298_v2.0.fa’) (Li 2018)
- Gene based alignments for grape vs peach generated with MCScan ‘python -m jcvi.compara.catalog ortholog grape peach --no_strip_names following’ using guide from https://github.com/tanghaibao/jcvi/wiki/MCscan-%28Python-version%29 (Tang et al. 2008)

The source code of JBrowse 2 is available at https://github.com/GMOD/jbrowse-components

Installation guides and documentation are available at https://jbrowse.org/jb2/docs A list of live demos associated with the figures in this paper is available at https://jbrowse.org/demos/paper2022/

## Acknowledgements

The expert interviews in the design sprints directly informed the software presented here. We especially wish to thank Jonathan Torchio, Jared Simpson, Fabien Lamaze, and Heather Gibling of OICR, Viswateja Nelakuditi and Hiromichi Suzuki of SickKids (The Hospital for Sick Children), Michael Schatz and Michael Alonge of Johns Hopkins University, and Xingang Wang of Cold Spring Harbor National Laboratory for their participation and invaluable feedback in design sprints. We also want to thank all contributors of code, feedback, bug and feature requests on the project’s GitHub pages.

## Author information

### Contributions

CD, GS, PX, TDJM, EH, JZ, CB, GH, AD, and RB formed the core software development team (led by RB) and worked on all areas of the project. AL contributed design elements and organized design sprints. EG worked on JBrowse Jupyter. MM, TF, and BB worked on text indexing. RH and SC worked on outreach. LDS and IHH led the project including preparation of manuscripts and grant proposals and overall direction.

## Ethics declarations

### Ethics approval and consent to participate

Not applicable.

### Consent for publication

Not applicable.

## References

“Apache License, Version 2.0.” n.d. Accessed April 15, 2022. https://opensource.org/licenses/Apache-2.0.

Buels, Robert, Eric Yao, Colin M. Diesh, Richard D. Hayes, Monica Munoz-Torres, Gregg Helt, David M. Goodstein, et al. 2016. “JBrowse: A Dynamic Web Platform for Genome Visualization and Analysis.” Genome Biology 17 (April): 66. https://doi.org/10.1186/s13059-016-0924-1.

Carver, Tim J., Kim M. Rutherford, Matthew Berriman, Marie-Adele Rajandream, Barclay G. Barrell, and Julian Parkhill. 2005. “ACT: The Artemis Comparison Tool.” Bioinformatics 21 (16): 3422–23. https://doi.org/10.1093/bioinformatics/bti553.

“Chain Format.” n.d. Accessed April 28, 2022. https://genome.ucsc.edu/goldenPath/help/chain.html.

De Jesus Martinez, Teresa, Elliot A. Hershberg, Emma Guo, Garrett J. Stevens, Colin Diesh, Peter Xie, Caroline Bridge, et al. 2022. “JBrowse Jupyter: A Python Interface to JBrowse 2.” bioRxiv. https://doi.org/10.1101/2022.05.11.491552.

Durand, Neva C., James T. Robinson, Muhammad S. Shamim, Ido Machol, Jill P. Mesirov, Eric S. Lander, and Erez Lieberman Aiden. 2016. “Juicebox Provides a Visualization System for Hi-C Contact Maps with Unlimited Zoom.” Cell Systems 3 (1): 99. https://doi.org/10.1016/j.cels.2015.07.012.

Durbin, Richard, and Jean Thierry-Mieg. 1994. “The ACEDB Genome Database.” In Computational Methods in Genome Research, edited by Sándor Suhai, 45–55. Boston, MA: Springer US. https://doi.org/10.1007/978-1-4615-2451-9_4.

“Electron.” n.d. Accessed April 27, 2022. https://www.electronjs.org/.

Espejo Valle-Inclan, Jose, Nicolle J. M. Besselink, Ewart de Bruijn, Daniel L. Cameron, Jana Ebler, Joachim Kutzera, Stef van Lieshout, et al. 2022. “A Multi-Platform Reference for Somatic Structural Variation Detection.” Cell Genomics 2 (6): 100139. https://doi.org/10.1016/j.xgen.2022.100139.

Gillespie, Marc, Bijay Jassal, Ralf Stephan, Marija Milacic, Karen Rothfels, Andrea Senff-Ribeiro, Johannes Griss, et al. 2021. “The Reactome Pathway Knowledgebase 2022.” Nucleic Acids Research 50 (D1): D687–92. https://doi.org/10.1093/nar/gkab1028.

Griffith, Malachi, Nicholas C. Spies, Kilannin Krysiak, Joshua F. McMichael, Adam C. Coffman, Arpad M. Danos, Benjamin J. Ainscough, et al. 2017. “CIViC Is a Community Knowledgebase for Expert Crowdsourcing the Clinical Interpretation of Variants in Cancer.” Nature Genetics 49 (2): 170–74. https://doi.org/10.1038/ng.3774.

Haas, Brian J., Alex Dobin, Nicolas Stransky, Bo Li, Xiao Yang, Timothy Tickle, Asma Bankapur, et al. 2017. “STAR-Fusion: Fast and Accurate Fusion Transcript Detection from RNA-Seq.” bioRxiv. https://doi.org/10.1101/120295..

Hershberg, Elliot A., Garrett Stevens, Colin Diesh, Peter Xie, Teresa De Jesus Martinez, Robert Buels, Lincoln Stein, and Ian Holmes. 2021. “JBrowseR: An R Interface to the JBrowse 2 Genome Browser.” Bioinformatics 37 (21): 3914–15. https://doi.org/10.1093/bioinformatics/btab459.

Hornik, Kurt. 2012. “The Comprehensive R Archive Network.” Wiley Interdisciplinary Reviews. Computational Statistics 4 (4): 394–98. https://doi.org/10.1002/wics.1212.

“HTS Format Specifications.” n.d. Accessed April 28, 2022. https://samtools.github.io/hts-specs/.

Jain, Chirag, Alexander Dilthey, Sergey Koren, Srinivas Aluru, and Adam M. Phillippy. 2018. “A Fast Approximate Algorithm for Mapping Long Reads to Large Reference Databases.” Journal of Computational Biology: A Journal of Computational Molecular Cell Biology 25 (7): 766–79. https://doi.org/10.1089/cmb.2018.0036.

“JavaScript With Syntax For Types.” n.d. Accessed April 15, 2022. https://www.typescriptlang.org/.

Kent, W. James, Charles W. Sugnet, Terrence S. Furey, Krishna M. Roskin, Tom H. Pringle, Alan M. Zahler, and David Haussler. 2002. “The Human Genome Browser at UCSC.” Genome Research 12 (6): 996–1006. https://doi.org/10.1101/gr.229102.

Krzywinski, Martin, Jacqueline Schein, İnanç Birol, Joseph Connors, Randy Gascoyne, Doug Horsman, Steven J. Jones, and Marco A. Marra. 2009. “Circos: An Information Aesthetic for Comparative Genomics.” Genome Research 19 (9): 1639–45. https://doi.org/10.1101/gr.092759.109.

Kurtz, Stefan, Adam Phillippy, Arthur L. Delcher, Michael Smoot, Martin Shumway, Corina Antonescu, and Steven L. Salzberg. 2004. “Versatile and Open Software for Comparing Large Genomes.” Genome Biology 5 (2): 1–9. https://doi.org/10.1186/gb-2004-5-2-r12.

Le Clainche Giorgio Malacrida Eléonore Durand Graziano Pesole Valérie Laucou Philippe Chatelet Didier Merdinoglu Massimo Delledonne Mario Pezzotti Alain Lecharny Claude Scarpelli François Artiguenave M. Enrico Pè Giorgio Valle Michele Morgante Michel Caboche Anne-Françoise Adam-Blondon Jean Weissenbach Francis Quétier & Patrick Wincker, Olivier Jaillon Jean-Marc Aury Benjamin Noel Alberto Policriti Christian Clepet Alberto Casagrande Nathalie Choisne Sébastien Aubourg Nicola Vitulo Claire Jubin Alessandro Vezzi Fabrice Legeai Philippe Hugueney Corinne Dasilva David Horner Erica Mica Delphine Jublot Julie Poulain Clémence Bruyère Alain Billault Béatrice Segurens Michel Gouyvenoux Edgardo Ugarte Federica Cattonaro Véronique Anthouard Virginie Vico Cristian Del Fabbro Michaël Alaux Gabriele Di Gaspero Vincent Dumas Nicoletta Felice Sophie Paillard Irena Juman Marco Moroldo Simone Scalabrin Aurélie Canaguier Isabelle. 2007. “The Grapevine Genome Sequence Suggests Ancestral Hexaploidization in Major Angiosperm Phyla.” Nature 449 (7161): 463–67. https://doi.org/10.1038/nature06148.

Li, Heng. 2018. “Minimap2: Pairwise Alignment for Nucleotide Sequences.” Bioinformatics 34 (18): 3094–3100. https://doi.org/10.1093/bioinformatics/bty191.

Loman, Nicholas J., Joshua Quick, and Jared T. Simpson. 2015. “A Complete Bacterial Genome Assembled de Novo Using Only Nanopore Sequencing Data.” Nature Methods 12 (8): 733–35. https://doi.org/10.1038/nmeth.3444.

Markham, John F., Satwica Yerneni, Georgina L. Ryland, Huei San Leong, Andrew Fellowes, Ella R. Thompson, Wasanthi De Silva, et al. 2019. “CNspector: A Web-Based Tool for Visualisation and Clinical Diagnosis of Copy Number Variation from next Generation Sequencing.” Scientific Reports 9 (1): 1–9. https://doi.org/10.1038/s41598-019-42858-8.

McKay, Sheldon J., Ismael A. Vergara, and Jason E. Stajich. 2010. “Using the Generic Synteny Browser (GBrowse_syn).” Current Protocols in Bioinformatics / Editoral Board, Andreas D. Baxevanis … [et Al.] Chapter 9 (September): Unit 9.12. https://doi.org/10.1002/0471250953.bi0912s31.

Nattestad, Maria, Robert Aboukhalil, Chen-Shan Chin, and Michael C. Schatz. 2020. “Ribbon: Intuitive Visualization for Complex Genomic Variation.” Bioinformatics 37 (3): 413–15. https://doi.org/10.1093/bioinformatics/btaa680.

Nattestad, Maria, Sara Goodwin, Karen Ng, Timour Baslan, Fritz J. Sedlazeck, Philipp Rescheneder, Tyler Garvin, et al. 2017. “Complex Rearrangements and Oncogene Amplifications Revealed by Long-Read DNA and RNA Sequencing of a Breast Cancer Cell Line.” bioRxiv. https://doi.org/10.1101/174938.

“Npm.” n.d. Accessed April 27, 2022. https://npmjs.com/.

Primrose: Predict 5mC in PacBio HiFi Reads. n.d. Github. Accessed April 28, 2022. https://github.com/PacificBiosciences/primrose.

Puppeteer: Headless Chrome Node.js API. n.d. Github. Accessed April 27, 2022. https://github.com/puppeteer/puppeteer.

“PyPI · the Python Package Index.” n.d. PyPI. Accessed April 27, 2022. https://pypi.org/.

Raney, Brian J., Timothy R. Dreszer, Galt P. Barber, Hiram Clawson, Pauline A. Fujita, Ting Wang, Ngan Nguyen, et al. 2013. “Track Data Hubs Enable Visualization of User-Defined Genome-Wide Annotations on the UCSC Genome Browser.” Bioinformatics 30 (7): 1003–5. https://doi.org/10.1093/bioinformatics/btt637.

“React.” n.d. Accessed April 15, 2022. https://reactjs.org/.

Robinson, James T., Helga Thorvaldsdóttir, Douglass Turner, and Jill P. Mesirov. 2020. “Igv.js: An Embeddable JavaScript Implementation of the Integrative Genomics Viewer (IGV).” bioRxiv. bioRxiv. https://doi.org/10.1101/2020.05.03.075499.

Robinson, James T., Helga Thorvaldsdóttir, Aaron M. Wenger, Ahmet Zehir, and Jill P. Mesirov. 2017. “Variant Review with the Integrative Genomics Viewer.” Cancer Research 77 (21): e31–34. https://doi.org/10.1158/0008-5472.CAN-17-0337.

Sedlazeck, Fritz J., Philipp Rescheneder, Moritz Smolka, Han Fang, Maria Nattestad, Arndt von Haeseler, and Michael C. Schatz. 2018. “Accurate Detection of Complex Structural Variations Using Single-Molecule Sequencing.” Nature Methods 15 (6): 461–68. https://doi.org/10.1038/s41592-018-0001-7.

Skinner, Mitchell E., Andrew V. Uzilov, Lincoln D. Stein, Christopher J. Mungall, and Ian H. Holmes. 2009. “JBrowse: A next-Generation Genome Browser.” Genome Research 19 (9): 1630–38. https://doi.org/10.1101/gr.094607.109.

Stein, Lincoln D., Christopher Mungall, Shengqiang Shu, Michael Caudy, Marco Mangone, Allen Day, Elizabeth Nickerson, et al. 2002. “The Generic Genome Browser: A Building Block for a Model Organism System Database.” Genome Research 12 (10): 1599–1610. https://doi.org/10.1101/gr.403602.

Tang, Haibao, John E. Bowers, Xiyin Wang, Ray Ming, Maqsudul Alam, and Andrew H. Paterson. 2008. “Synteny and Collinearity in Plant Genomes.” Science. https://doi.org/10.1126/science.1153917.

The Ensembl Core Team. n.d. “The Track Hub Registry.” Accessed April 27, 2022. https://www.trackhubregistry.org/.

“The Variant Call Format Specification - VCFv4.3 and BCFv2.2.” 2022. April 19, 2022. https://samtools.github.io/hts-specs/VCFv4.3.pdf.

“Trix Indices.” n.d. Accessed April 15, 2022. https://genome.ucsc.edu/goldenPath/help/trix.html.

“Using Web Workers.” n.d. Accessed April 27, 2022. https://developer.mozilla.org/en-US/docs/Web/API/Web_Workers_API/Using_web_workers.

Veltri, Daniel, Martha Malapi Wight, and Jo Anne Crouch. 2016. “SimpleSynteny: A Web-Based Tool for Visualization of Microsynteny across Multiple Species.” Nucleic Acids Research 44 (W1): W41–45. https://doi.org/10.1093/nar/gkw330.

Verde, Ignazio, Albert G. Abbott, Simone Scalabrin, Sook Jung, Shengqiang Shu, Fabio Marroni, Tatyana Zhebentyayeva, et al. 2013. “The High-Quality Draft Genome of Peach (Prunus Persica) Identifies Unique Patterns of Genetic Diversity, Domestication and Genome Evolution.” Nature Genetics 45 (5): 487–94. https://doi.org/10.1038/ng.2586.

Weitz, Eric. n.d. Ideogram: Chromosome Visualization for the Web. Github. Accessed April 27, 2022. https://github.com/eweitz/ideogram.

“Welcome to MobX-State-Tree!” n.d. Accessed April 15, 2022. https://mobx-state-tree.js.org//.

Xin, Jiwen, Adam Mark, Cyrus Afrasiabi, Ginger Tsueng, Moritz Juchler, Nikhil Gopal, Gregory S. Stupp, et al. 2016. “High-Performance Web Services for Querying Gene and Variant Annotation.” Genome Biology 17 (1): 91. https://doi.org/10.1186/s13059-016-0953-9.

Yokoyama, T. T., and M. Kasahara. 2020. “Visualization Tools for Human Structural Variations Identified by Whole-Genome Sequencing.” Journal of Human Genetics 65 (1). https://doi.org/10.1038/s10038-019-0687-0.

