## Supplementary info for "JBrowse 2: A modular genome browser with views of synteny and structural variation"

#### Supplementary Information

#### Performance profiling

We looked at the time to render a single track on 5kb, 10kb, and 19kb region sizes, to show patterns where larger regions involve fetching and drawing more data. Above 19kb, the browsers (by default) decline to display information, instead suggesting that the user “Zoom in to see more info”.

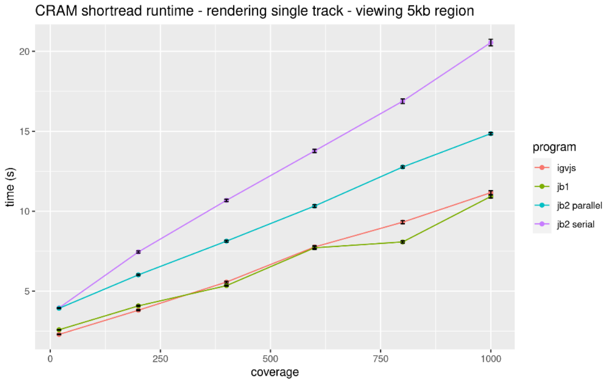


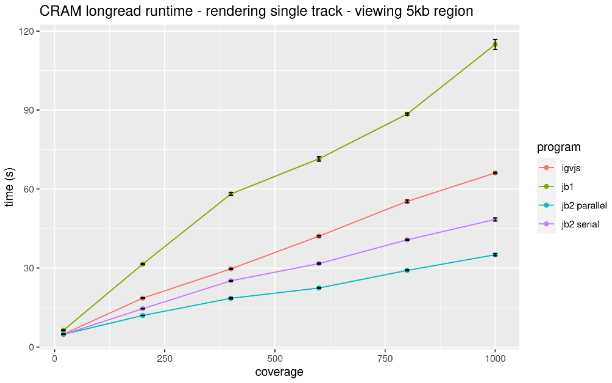


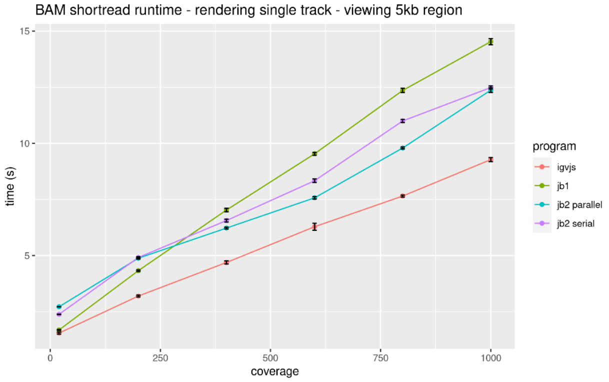


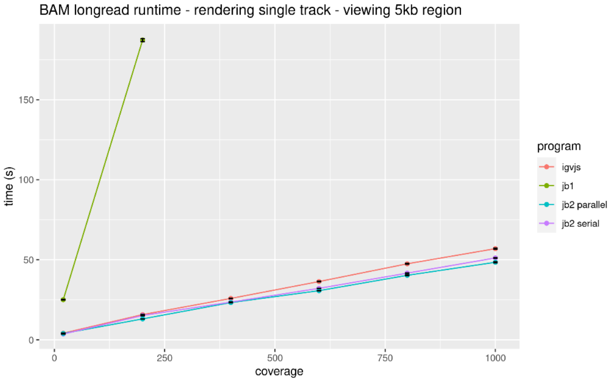

CRAM shortread runtime - rendering single track - viewing 10kb region

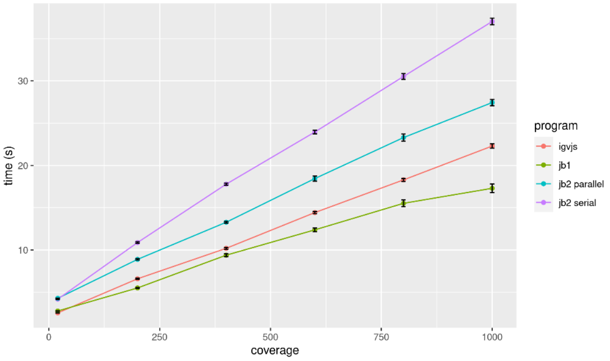


CRAM longread runtime - rendering single track - viewing 10kb region

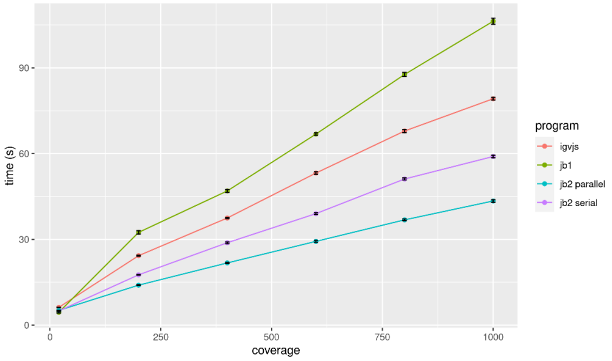


BAM shortread runtime - rendering single track - viewing 10kb region

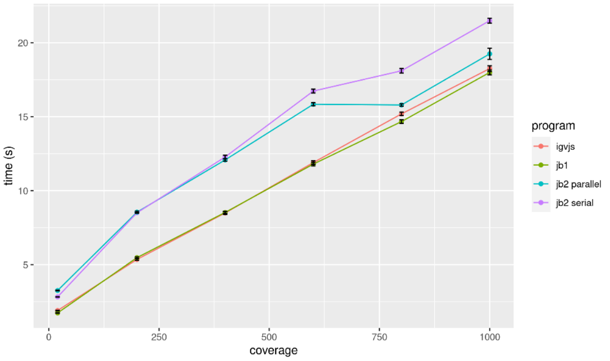


BAM longread runtime - rendering single track - viewing 10kb region

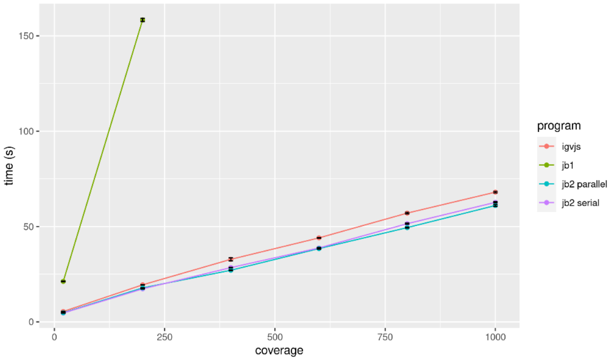


CRAM shortread runtime - rendering single track - viewing 19kb region

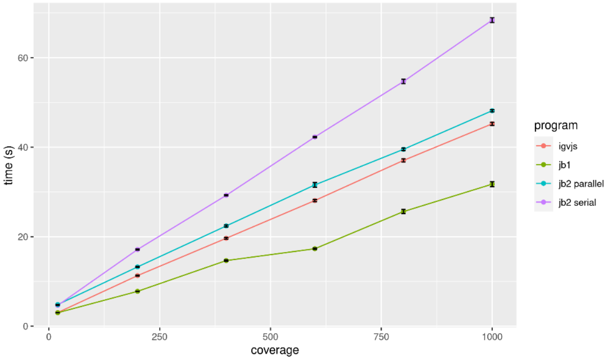


CRAM longread runtime - rendering single track - viewing 19kb region

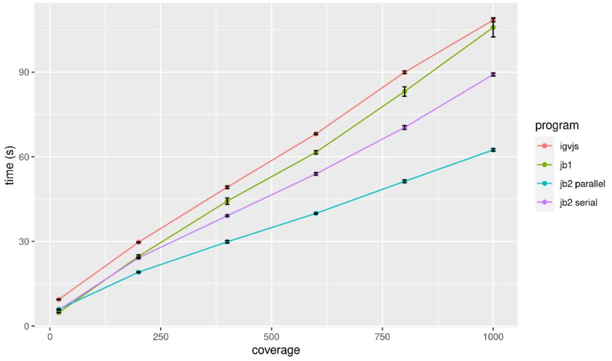


BAM shortread runtime - rendering single track - viewing 19kb region

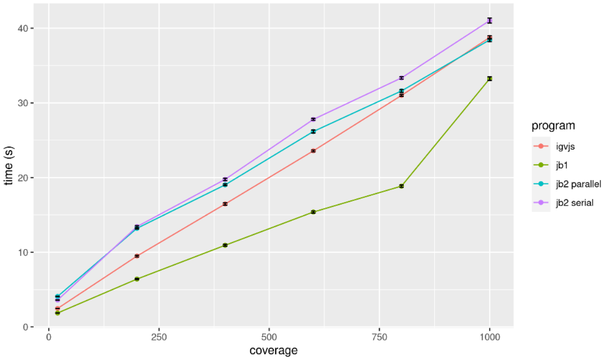


BAM longread runtime - rendering single track - viewing 19kb region

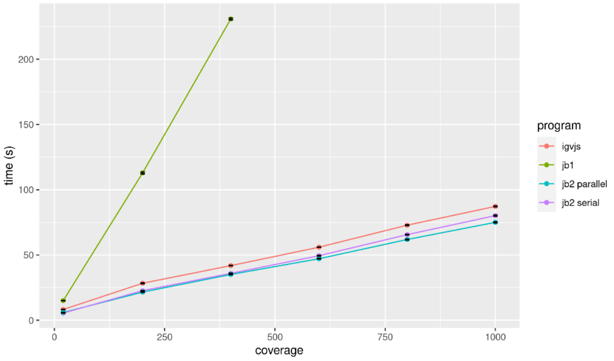

Figure S1: The performance of JBrowse 2 Web (v1.7.7), JBrowse 2 Embedded (v1.7.7), JBrowse 1 (v1.16.11) and igv.js (v2.12.1) displaying a single BAM or CRAM track at varying coverages (see section "Performance and Scalability for benchmark details"). The error bars are indicated in black on each measurement. Incomplete lines

### Calculation of empirical distribution of time between frames

If the frame lengths are $\{t_i: 1 \le i \le N\}$ then the total benchmark duration is $T = \sum_{i=1}^{N} t_i$

Let $U$ be a random instant in time, uniformly distributed over the total benchmark duration.
Let $W$ be the wait time from $U$ until the start of the next frame.

Then $W$ is a natural measure of "responsiveness", since it represents the (random) delay from when the user tries to perform an action, to when the app responds.
The probability that $U$ is in the $i$'th time interval is $\frac{t_i}{T}$; conditional on this, $W$ is uniformly distributed over $[0, t_i]$.

Using this, the expectation and variance of $W$ are (although we don't need these directly)

$$E[W] = \sum_{i=1}^{N} \frac{t_i^2}{2T}$$

$$V[W] = \left(\sum_{i=1}^{N} \frac{t_i^3}{3T}\right) - E[W]^2$$

The cumulative distribution function (cdf) of $W$ is

$$P(W \le X) \equiv C(X) = \frac{1}{T}\sum_{i=1}^{T} min\left(t_i, X\right)$$

We want to invert $C$, i.e. find $X = C^{-1}(Q)$ such that $C(X) = Q$.

Let $L(X) = \left|\{t_i \le X\}\right|$ be the number of frames that render in time $X$ or less.
Let $\{u_i\}$ be the set of frame times $\{t_i\}$ sorted in ascending order, so $u_{L(X)} \le X < u_{L(X)+1}$.
For convenience define $u_0 = 0$. Then we can rewrite the cdf of $W$ as

$C(X) = \frac{1}{T}\left(X \times (N - L(X)) + S_{L(X)}\right)$ where $S_k = \sum_{i=0}^{k} u_i$.

Note that $C(X)$ is piecewise linear with joins at $\{u_i\}$. Let $F(Q) = max\{k: 0 \le k \le N, C(u_k) \le Q\}$ be the index of the last join at or before the point where $C(X)$ reaches $Q$. It follows that $u_{F(Q)} \le C^{-1}(Q) < u_{F(Q)+1}$ and $L(C^{-1}(Q)) = F(Q)$.
Thus we can again rewrite the cdf of $W$ as
$Q = \frac{1}{T}(X \times (N - F(Q)) + S_{F(Q)})$
which we can rearrange for $X$ to give the required formula for the quartile points of the cdf as follows

$$C^{-1}(Q) = \frac{QT - S_{F(Q)}}{N - F(Q)}$$

### Data parser libraries

Table S1 lists data parser and accessory libraries created during JBrowse 2 development.

| **NPM package name** | **Purpose** |
|---|---|
| @gmod/bam | Read BAM files |
| @gmod/bbi | Read UCSC BigWig and BigBed files |
| @gmod/bed | Read BED files with or without autoSql schemas |
| @gmod/bgzf-filehandle | Read block-gzipped (BGZip) data files that support random access |
| @gmod/cram | Read CRAM files (Buels et al. 2019) |
| @gmod/faidx | Create FASTA indexes |
| @gmod/gff | Read GFF3 files |
| @gmod/gtf | Read GTF feature files |
| @gmod/indexedfasta | Read indexed FASTA or bgzip indexed fasta files |
| @gmod/nclist | Read JBrowse 1 NCList track data |
| @gmod/tabix | Read Tabix-indexed files |
| @gmod/trix | Read UCSC trix indexes |
| @gmod/twobit | Read UCSC .2bit files |

| | |
|---|---|
| @gmod/ucsc-hub | Read UCSC track hub definitions |
| @gmod/vcf | Read VCF files |
| abortable-promise-cache | Asynchronous cache that can cancel long-running operations |
| generic-filehandle | Transparently read from either network resources or local files |
| ixixx | Create UCSC trix indexes |

Table S1. Listing of selected NPM modules produced by JBrowse 2 development.
